# Ultra-fast Prediction of Somatic Structural Variations by Reduced Read Mapping via Pan-Genome *k*-mer Sets

**DOI:** 10.1101/2020.10.25.354456

**Authors:** Min-Hak Choi, Jang-il Sohn, Dohun Yi, A Vipin Menon, Yeon Jeong Kim, Sungkyu Kyung, Seung-Ho Shin, Byunggook Na, Je-Gun Joung, Sungro Yoon, Youngil Koh, Daehyun Baek, Tae-Min Kim, Jin-Wu Nam

**Affiliations:** Department of Life Science, Hanyang University, Seoul 04763, Republic of Korea; Research Institute for Convergence of Basic Sciences, Hanyang University, Seoul 04763, Republic of Korea; Research Institute for Natural Sciences, Hanyang University, Seoul 04763, Republic of Korea; Samsung Genome Institute, Samsung Medical Center, 81 Irwon-ro, Gangnam-gu, Seoul, 06351, Republic of Korea; GENINUS Inc, Seoul 05836, Republic of Korea; Department of Electrical and Computer Engineering, Seoul National University, 08826, Republic of Korea; College of Medicine, Seoul National University, Seoul 03080, Republic of Korea; School of Biological Sciences, Seoul National University, Seoul 08826, Republic of Korea; Department of Medical Informatics and Cancer Research Institute, College of Medicine, The Catholic University of Korea, Seoul 06591, Republic of Korea

**Keywords:** Somatic structural variation, whole genome sequencing, cancer panel sequencing, fusion gene, pan-genome *k*-mer set, reduced mapping

## Abstract

Genome rearrangements often result in copy number alterations of cancer-related genes and cause the formation of cancer-related fusion genes. Current structural variation (SV) callers, however, still produce massive numbers of false positives (FPs) and require high computational costs. Here, we introduce an ultra-fast and high-performing somatic SV detector, called ETCHING, that significantly reduces the mapping cost by filtering reads matched to pan-genome and normal *k*-mer sets. To reduce the number of FPs, ETCHING takes advantage of a Random Forest classifier that utilizes six breakend-related features. We systematically benchmarked ETCHING with other SV callers on reference SV materials, validated SV biomarkers, tumor and matched-normal whole genomes, and tumor-only targeted sequencing datasets. For all datasets, our SV caller was much faster (≥15X) than other tools without compromising performance or memory use. Our approach would provide not only the fastest method for largescale genome projects but also an accurate clinically practical means for real-time precision medicine.

## Introduction

Chromosomal rearrangements in coding regions and regulatory non-coding elements often cause malignancy of somatic cells. Although structural variations (SVs) occur much less frequently than single nucleotide variations (SNVs), the SVs often have a greater impact on cellular functions and gene expression ^1^. In particular, large SVs (>1Kbp), which include large insertions (INSs), deletions (DELs), inversions (INVs), duplications (DUPs), and translocations (TRAs), are more often associated with gain- and/or loss-of-function of cancer-related genes and druggable target genes for cancer treatments than are SNVs ^2–7^. For instance, *ERBB2* amplification in breast cancers (BRCAs) ^8, 9^, *EML4*-*ALK* fusion in lung cancer ^10^, and *BCR*-*ABL* fusion in chronic myeloid leukemia ^11^ are well-known SV-driven cancer drivers and actionable targets for cancer treatments. Hence, the rapid detection of cancer-related SVs is indispensable for companion diagnostics and targeted cancer therapy.

So far, a handful of SV callers have been introduced to find germline and somatic SVs in normal and/or tumor samples by using a read-based approach –– read-depth ^12^, discordant read-pairs ^13^, soft-clipped reads ^14–16^, and their combinations ^17–21^ –– or by using a *k*-mer-based approach ^22^. Some of them utilize local assembly of reads ^13, 20–22^ to precisely detect breakpoints (BPs) and SV types. Regardless of the approach, all current SV callers require genome mapping of all input reads. Although the mapping process is an indispensable step for the confident identification of SVs, it consumes most of the computing time in processing massive whole genome sequencing (WGS) data. For instance, the genome mapping of 30X WGS data from a cancer patient takes ~300 hours with a single thread on a high-performing computer, resulting in delayed diagnosis. Furthermore, SV studies for largescale WGS projects, such as those undertaken by the Pan-Cancer Analysis of Whole Genomes (PCAWG) ^23^ and the Genome Aggregation Database (gnomAD) ^24^ consortiums, would be only doable by institutes with access to a giant computing facility or expensive cloud computing services.

The majority of sequenced reads are reference reads (perfectly matched to the reference genome), which could be dispensable for SV calling. Mapping the reference reads consumes expensive computing time. It may also increase background noise resulting from imprecise and ambiguous alignments of the reads (mainly due to repeats or low-complexity regions) or from unresolved misassemblies of the reference ^25, 26^. Thus, only mapping informative (non-reference) reads to detect SVs would both reduce computing time and increase accuracy.

In general, somatic SV callers use a case-control design that compares tumor (case) SVs with those of matched normals (control) to detect somatic (case-specific) SVs. The absence of matched normal samples may lead to either a failure of SV-calling or a high FP rate spawned by germline SVs. In particular, cancer panel sequencing is frequently carried out using only tumor samples. Using the pan-genome sequences containing all non-medical variations instead of a matched normal sample would help to enhance the accuracy of SV calling in this situation.

In this study, we developed ETCHING, an ultra-fast SV detection method. Our approach significantly reduces the number of reads to be mapped by excluding those from the reference and/or pan-genome *k*-mer (PGK) set. This new strategy drastically reduces running time (it is at least ~15 times faster than other methods) without compromising performance by taking advantage of machine-learning-based classification to remove FP SVs further. ETCHING displays either comparable to or better accuracy than other state-of-the-art SV detection tools on benchmarking whole genome and panel sequencing datasets as well as reference materials.

## Results

### Fast prediction of somatic SVs

We report the development of ETCHING (Efficient deTection of CHromosomal rearrangements and fusIoN Genes) – a fast computational SV caller that comprises four stepwise modules: Filter, Caller, Sorter, and Fusion-identifier (Fig. 1a; Supplementary Fig. 1; see Methods for more details). The Filter module uses one of three different filters: a Pan-Genome *k*-mer (PGK) filter that excludes tumor reads in which all *k*-mers are present in PGK, a Normal filter that removes those reads in which all *k*-mers come from normal reads (not using reference genomes), or a combined (PGKN) filter (Fig. 1b). PGK is a unique set of 31-mers from 10 human genome assemblies and nonpathogenic single nucleotide polymorphisms (SNPs) from dbSNP (~3.9 × 10^9^ *k*-mers; Supplementary Fig. 2; Supplementary Table 1). This module allows us to collect tumor-specific (TS) reads by filtering reference reads, those with germline variations, and those matched to normal reads. We used The Cancer Genome Atlas (TCGA) BRCA WGS data used in a previous SV study ^27^ for checking the Filter module. Of the BRCA samples, 31 and 9 were selected for training and hold-out test, respectively, by random selection (Methods; Supplementary Table 2). For the hold-out test dataset, the Filter module excluded about 96.2% of the reads by PGK, 99.2% by Normal, and 99.4% by PGKN (Fig. 1c). The remaining TS reads clearly present BPs with a sharp decay of read-depth in somatic DEL, DUP, INV, and TRA examples, reminiscent of the chemical etching process (Fig. 1d). This filtration method significantly shortened the mapping process. The mapping time for TS reads from the nine hold-out BRCA WGS datasets with varied coverages (33–68X and 27–56X in tumor and normal samples, respectively) was approximately 300 times faster than that for all reads (Unfiltered) using BWA-MEM ^28^ (Fig. 1e).

**Fig. 1.**
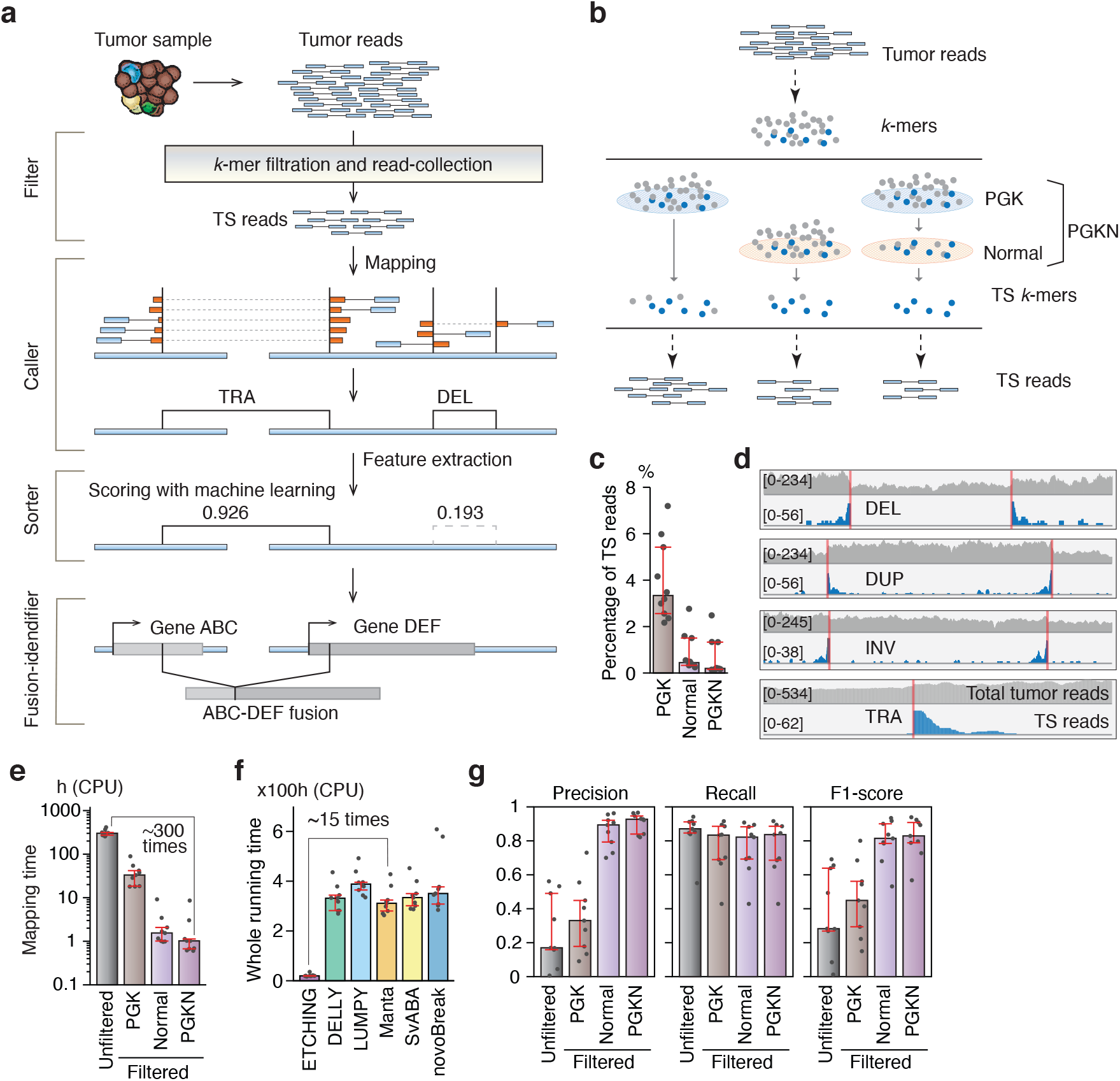
A schematic overview of ETCHING. **a.** A schematic showing the flow through the ETCHING process, which comprises four stepwise modules (Filter, Caller, Sorter, and Fusion-identifier). **b.** The Filter module collects TS reads containing at least one TS *k*-mer not present in the *k*-mer sets (PGK, Normal, or PGKN). **c.** The percentage of TS reads that pass through the PGK, Normal, and PGKN filters. **d.** The mapping patterns of the total tumor reads (unfiltered, gray) and TS reads (filtered, blue) are shown for representative DEL, DUP, INV, and TRA loci via Integrative Genome Viewer. **e.** The mapping times (CPU times) required for the total tumor reads (Unfiltered) and TS reads filtered by PGK, Normal, and PGKN using BWA-MEM. **f.** The total running time (CPU time) of the SV callers. **g.** The precision, recall, and F1-scores of ETCHING with total tumor (Unfiltered) and TS reads collected by PGK, Normal, and PGKN. (**c,e-g**) The analyses were done with nine BRCA WGS datasets. The error bars indicate the first to third quartile range, and the height of the boxes indicate median values.

After mapping TS reads to the reference genome (hg19), the Caller module collects simple-clipped reads to find initial BPs (Supplementary Fig. 3a) and then defines breakends (BNDs) for BP pairs by considering the clipped direction (Supplementary Fig. 3b). The identified BNDs were then assigned to an SV type, such as DEL, DUP, INV, and TRA, according to their position and the clipped direction (Supplementary Fig. 3c; Methods). Next, the Sorter module predicts a confidence score for each SV call using machine learning models pre-trained over the 31 training datasets (Methods). Because there is no ground truth for the TCGA dataset, we instead used a silver standard set of SVs, simultaneously detected by multiple SV callers, during training and evaluation (Methods). Random Forest ^29^ (RF)-based sorter was chosen as our default SV sorter module (Methods; Supplementary Fig. 4). In the last step, with the predicted SVs, the Fusion-identifier module predicts fusion-gene (FG) candidates (Methods).

We compared the running time of ETCHING with those of other SV callers over the hold-out test dataset. The CPU time (running time converted in a single thread) for the entire SV prediction process of ETCHING was at least 15 times less than those of the other SV callers (Fig. 1f). In real (wall-clock) time, ETCHING took 2.2 hours on average, meaning that it was at least 6.6 times faster than the second-fastest caller (Manta), on 30 threads (Supplementary Fig. 5). The ETCHING process not only reduced the running time but also increased the precision of the SV prediction (Fig. 1g). The PGK, Normal, and PGKN filters gradually reduced the number of FP reads with little compromise of the true positive (TP) rates, resulting in better performances (F1-score) with the filters on BRCA WGS and HCC1395 cell line WGS datasets (Fig. 1g; Supplementary Fig. 6). Taken together, these results suggest that ETCHING provides high-performance SV prediction at a faster rate than other SV callers.

### ETCHING displays robust performance

We next sought to systematically benchmark the performance of ETCHING against the performances of the read-based callers DELLY, LUMPY, Manta, and SvABA, as well as that of a *k*-mer-based caller, novoBreak, over WGS data from the HCC1395 cancer cell line (50X) and its matched normal cell line, HCC1395 BL (30X). Because the HCC1395 dataset also lacks ground-truth SVs, we again used the approach of employing silver standard SVs identified by multiple callers, mentioned above. The precision-recall (PR) curves over varying parameters showed that ETCHING performed more robustly than the other callers, particularly for precision (Fig. 2a). We obtained optimal cutoffs, which were used in the following benchmarking analyses for fair comparisons (Fig. 2a, red indicator; Methods).

**Fig. 2.**
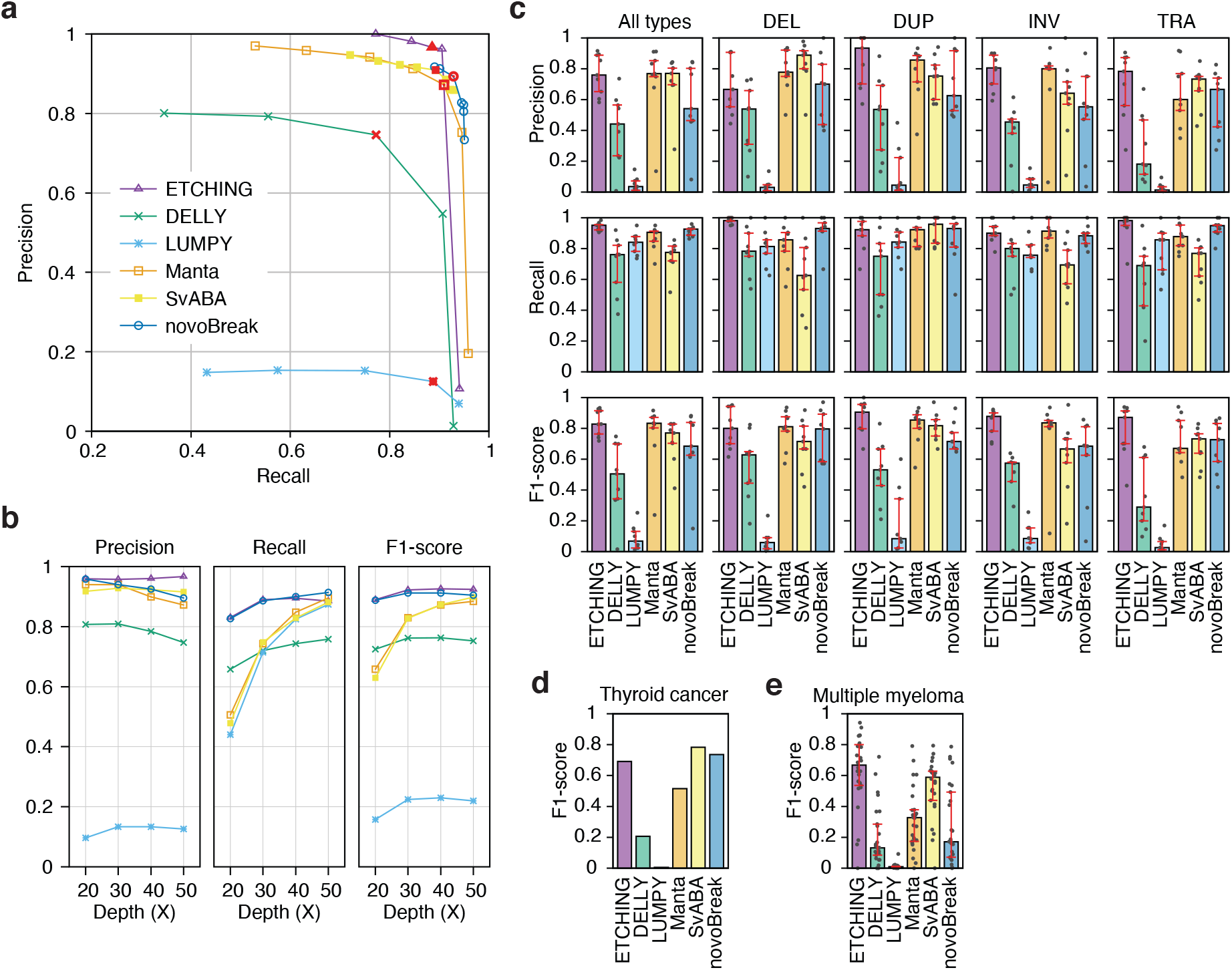
Performances of ETCHING and benchmarking SV callers. **a.** PR curves of ETCHING and benchmarking tools on the HCC1395 dataset. The red symbols indicate the points corresponding to optimal parameters. **b.** Precision, recall, and F1-scores of ETCHING and benchmarking tools over sub-sampled data with different sequencing depths from the HCC1395 tumor sample. **c.** Precision, recall, and F1-scores of ETCHING and benchmarking tools for all types of SVs over nine hold-out test datasets of TCGA BRCA samples. Each dot denotes the performance of each tool on a sample. The height of the bar plots indicates the median performance of each tool on nine samples, and the red error bars are the first and third quartiles. **d.** The performances of ETCHING and benchmarking tools on four TCGA THCA samples. Because there were only a few true SVs from each of the samples, we combined them as one value. **e.** The performances of ETCHING and benchmarking tools on MM samples. **d,e**. Otherwise, as in (**c**).

Because the performances of SV callers tend to be affected by the read-depth ^30^, we then examined the robustness of the SV callers over varying read-depths. For this comparison, we randomly subsampled 40% (20X), 60% (30X), and 80% (40X) of the reads from the HCC1395 cancer line (50X) while keeping the depth of the normal reads fixed, and then performed benchmarking analyses with the optimal cutoffs (Fig. 2b). ETCHING displayed a robust performance, regardless of the read-depth, and showed a slightly increased precision as the read-depth became higher. In contrast, Manta and SvABA presented lower recall rates at low read-depths but performances that were comparable to that of ETCHING at 50X.

To compare the performance of ETCHING on primary tumor samples with varying read-depth and tumor purity with those of the other tools, the nine hold-out BRCA samples were again used as the benchmarking dataset (Supplementary Table 2). In this analysis, ETCHING showed results that were superior or comparable to those of other tools, regardless of the SV type (Fig. 2c). Notably, ETCHING robustly predicted all SV types while displaying high F1-scores across samples, compared to other tools. We also benchmarked the SV callers over 33 true SVs from four thyroid cancer (THCA) samples of TCGA as an independent evaluation dataset. The performance of ETCHING was comparable to those of SvABA and novoBreak in terms of the F1-scores (Fig. 2d; Supplementary Fig. 7; Supplementary Table 3).

Because the silver standard set of SVs could still include FPs, we selected high-quality (HQ) SVs with depth-difference and connect-pair scores for DEL/DUP and INV/TRA, respectively (Supplementary Fig. 8; see Supplementary Note for details). With HQ SVs, ETCHING still displayed an accuracy that was comparable or superior to that of the other tools (Supplementary Fig. 9).

### SV prediction of experimentally validated targets

For experimental validation of the SV callers, we newly sequenced the whole genomes of 26 multiple myeloma (MM) samples with matched normal samples (Supplementary Table 4). We first benchmarked the SV callers using the MM samples, and found that ETCHING outperformed the others over a silver standard set of all SV types (Fig. 2e; Supplementary Fig. 9 and 10). Notably, its performance exceeded that of another *k*-mer-based caller, novoBreak, which showed a lower precision, particularly for INV and TRA types.

We then evaluated all of the SV callers using known clinical SV biomarkers of MM, such as DELs (in 1q25, *p16, RB1*, and *TP53*) and IGH rearrangements (including DELs and TRAs)(Fig. 3a) ^31^. Fluorescence *in situ* hybridization (FISH) and karyotype were first examined on the SV biomarkers (Supplementary Table 5). However, because FISH probe sets (Supplementary Table 6) of the SV biomarkers cannot discern focal deletion/duplication from (partial) aneuploidy, the true set of biomarker SVs were selected through a manual curation by considering read-depth changes and unbalanced minor allele frequency (Supplementary Fig. 11a,b) as well as discordant paired-reads in tumor and normal samples for each patient (Supplementary Fig. 11c–e; Methods). The SV set supported by FISH and/or karyotype (excluding aneuploidy) were well overlapped with manually curated SV biomarkers (Fig. 3b). We accordingly benchmarked ETCHING and other SV callers with the manually curated SV biomarkers as a true set. The receiver operating characteristics (ROC) showed that ETCHING displays comparable or slightly better performances than other callers in the SV biomarker level (Fig. 3c). Of 23 curated biomarkers, ETCHING detected 19. When breaking down the *IGH* rearrangements into SV level, known MM target genes, *FGFR3*, *IL6ST*, *CCND3*, *CCND1*, and *IGLL5* were detected as translocation partners by manual curation (Fig. 3b middle; Supplementary Table 7). Of the 38 SVs, ETCHING detected 17 SVs but missed seven including a *p16* DEL (MM17), three *IGH* DELs (MM10, 12, and 18), and three *IGH* TRAs (MM1, 11, and 14) (Fig. 3b).

**Fig. 3.**
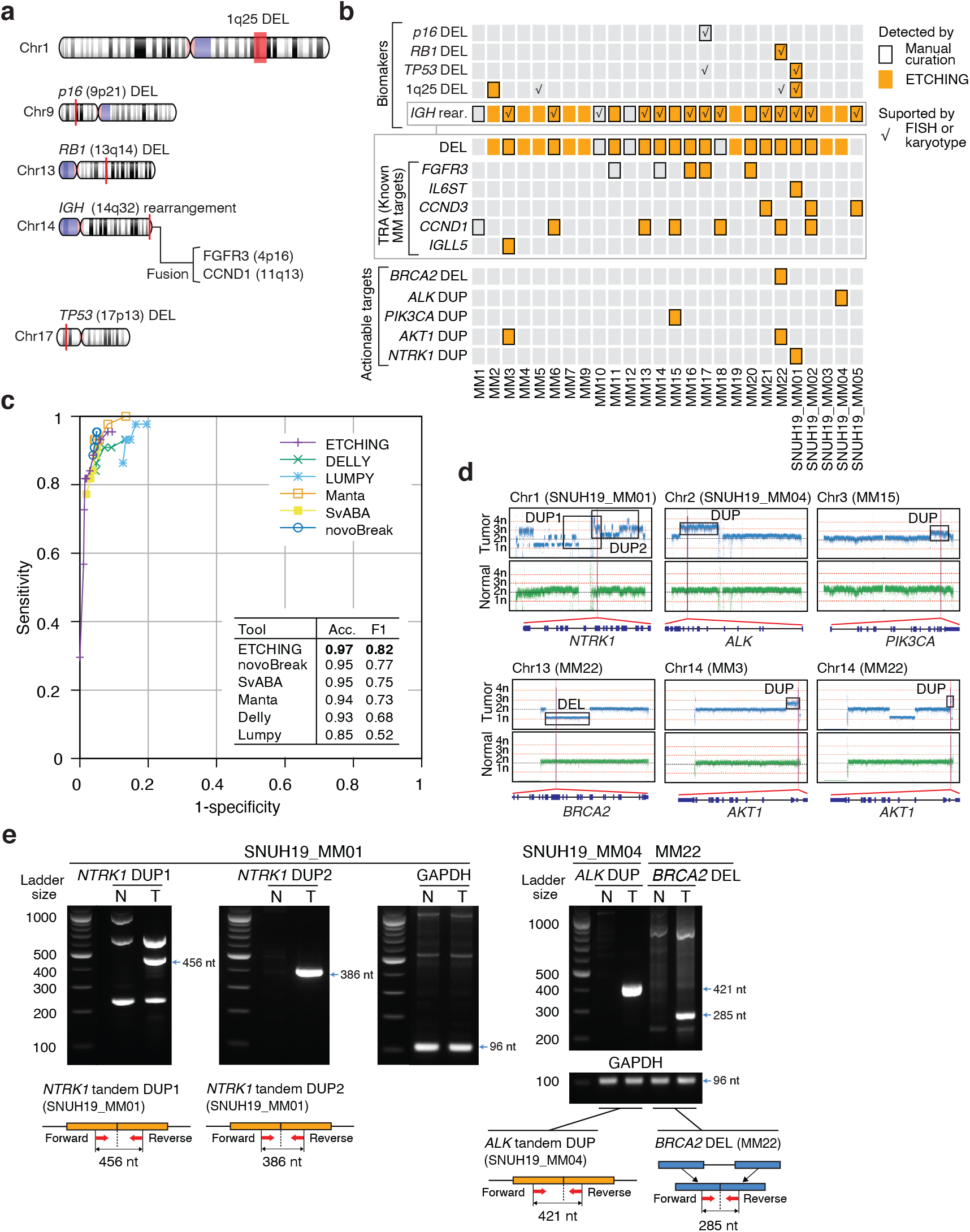
Prediction of SVs and FGs by SV callers using MM samples containing known clinical biomarkers and actionable SV targets. **a.** Known clinical SV and FG biomarkers (also known as clinical targets) of MM. The type of SV of known clinical biomarkers are indicated on the appropriate chromosomes. **b.** Summary of manually curated, experiment-supported, and ETCHING-detected SV biomarkers, known MM targets, and actionable targets from OncoKB (tier1, 2, and 3). **c.** ROC curves of ETCHING and benchmarking tools are shown along with accuracies (acc) and F1 scores as an inset. The accuracies and F1 scores were calculated on optimal parameters. **d**. The read-depth landscapes for chromosomes in which clinical biomarkers and targets were found. **e**. Experimental validation of three predicted actionable SV targets by PCR. The blue arrows indicate the expected sizes of the PCR amplicons in the gel images. ‘N’ indicates the normal sample and ‘T’ indicates the tumor sample. (bottom) The dotted lines indicate the junctions formed from tandem DUPs and DELs. The red arrows are the forward and reverse PCR primers.

We further searched for SVs related to actionable (cancer-druggable and clinically verified) targets from the OncoKB database ^32^. ETCHING detected five actionable SV targets – *BRCA2* DEL (MM22), *ALK* DUP (SNUH19_MM04), *PIK3CA* DUP (MM15), *AKT1* DUP (MM3), and *NTRK1* DUP (SNUH19_MM01) (Fig. 3b,d). Of the five predicted targets, three targets (excluding those from the MM3 and MM15 patients, which lack tumor DNA quantities) were verified by targeted PCR (Fig. 3e). The PCR products expected after amplification of *ALK* DUP (SNUH19_MM04), *BRCA2* DEL (MM22), and *NTRK1* DUPs (SNUH19_MM01) were observed in tumor but not in normal samples, indicating that the SV targets are true cases.

### SV and FG prediction in cancer panel sequencing

Targeted gene panel sequencing is more relevant than WGS for clinical applications, and clinical laboratories daily produce panel sequencing data with the aim of finding actionable target variations, SNVs, SVs, and FGs. Targeted gene panel sequencing is often applied to detect low-frequency alterations such as somatic SNVs or FGs in cell-free DNA from cancer patients. To test the effectiveness of the SV callers in such clinical situations, we analyzed 56 targeted gene panel sequencing data derived from three types of cell-free DNA (cfDNA) reference material (Methods): Complete Reference (CR), Complete Mutation Mix (CMM), and Mutation Mix v2 (MMv2). Each type contains two or three synthetic FGs with low mutant allele ratios (0.5–5%) and wild-type (WT) alleles from a cell line, GM24385.

Because cancer panel sequencing approaches generally lack matched normal data, ETCHING was first set to use a PGK filter to extract TS reads for SV prediction. Other benchmarking tools, with the exception of novoBreak, also predict SVs in the absence of normal data. novoBreak, given its requirement for normal data, used simulated data from the hg19 reference genome (Methods). Note that we ran all tools with default parameters that display a better recall rate for panel sequencing data.

This analysis showed that, along with LUMPY and DELLY, ETCHING is one of the top callers in terms of recall over such low mutant allele frequencies (Fig. 4a,b), while showing a moderate level of additional calls in targeted regions (Fig. 4c). Additional calls could be either FP calls or germline SVs from the WT sample. Compared to other tools, ETCHING barely predicted additional calls in non-target regions, indicating a relatively low frequency of FP calls (Fig. 4c, gray). Because the reference materials include WT data that lack mutant alleles, the benchmarking analyses of SV prediction were also performed using the WT data as the normal sample. ETCHING was then set to use a PGKN filter. Unlike the other tools, ETCHING and LUMPY maintained high recall rates (Supplementary Fig. 12) compared to the results obtained without WT data (Fig. 4a,b). This result indicates that ETCHING and LUMPY can effectively remove FPs without compromising the recall rate for targeted gene panel sequencing data, regardless of the presence of matched normal data. Since BreaKmer^33^ is specialized for targeted sequencing data, we also tested it on the same dataset. However, BreaKmer failed to report any result (Methods).

**Fig. 4.**
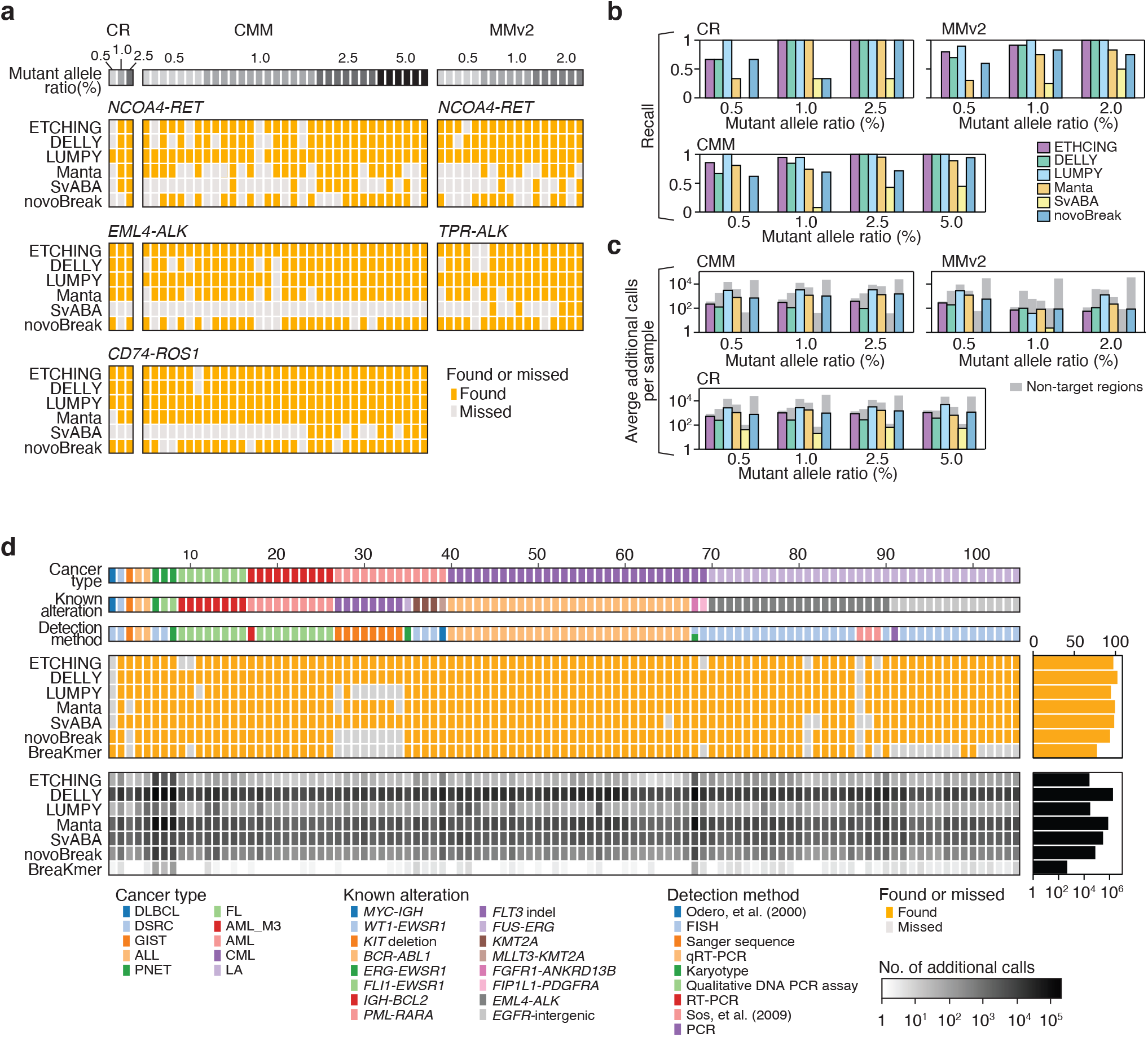
SV and FG predictions on targeted gene panel sequencing data. **a.** The TP calls (labeled as ‘Found’ in orange) and false negatives (labeled as ‘Missed’ in gray) of SV callers for cfDNA reference materials – CR, CMM, and MMv2 – with different mutant allele ratios (0.5 to 5.0%; gray to black). CR and CMM include *NCOA4-RET*, *EML4-ALK*, and *CD74-ROS1* FGs, and MMv2 includes *NCOA4-RET* and *TPR-ALK* FGs. **b**. The recall rates of benchmarking SV callers on the reference materials across different mutant allele ratios. **c.** The additional calls in target regions (colors) and non-target regions (gray). **d**. The heatmap summarizes the TPs (labeled as ‘Found’ in orange), false negatives (labeled as ‘Missed’ in gray), and additional calls for 105 cancer panel sequencing datasets. The panels on the right show the total number of TPs and additional calls. The white-to-black gradient indicates the number of additional calls on each SV caller. The color-coded charts (top) indicate cancer types, known alterations, and detection methods. Abbreviations: Diffuse large B-cell lymphoma (DLBCL), desmoplastic small round cell tumor (DSRC), gastrointestinal stromal tumor (GIST), acute lymphoblastic leukemia (ALL), primitive neuroectodermal tumor (PNET), follicular B-cell lymphoma (FL), acute myeloid leukemia (AML), chronic myelogenous leukemia (CML), and lung adenocarcinoma (LA).

Then, on the cancer panel sequencing data from formalin-fixed paraffin-embedded (FFPE) and frozen tissues from a previous study of BreaKmer ^33^, we evaluated the performances of ETCHING and other tools including BreaKmer. The data consists of 105 replicates from 37 samples of different types of cancers (Supplementary Table 8). Because the data included tumor samples without matched normal samples, ETCHING utilized the PGK, rather than the PGKN, filter for this prediction as above. All settings for the other tools were the same as were used for the reference materials. Since the data contains small variants in *FLT3* and *KIT*, we included small variations in this analysis (Methods). We first ran BreaKmer and compared its results to those of the previous study ^33^. BreaKmer was still very specific, giving only 479 additional calls across all 105 cases. However, it showed a lower recall rate (78 out of 105) than that they reported. It is possibly due to the lack of bait information or using a different version of BreaKmer (Methods). Other tools showed comparable recall rates (94 to 103 out of 105). Although DELLY showed the most sensitive performance, it was at the cost of massive additional calls (about one million). ETCHING found 98 true variants, and its number of additional calls was the lowest level except BreaKmer.

ETCHING was one of three tools that were able to detect all eight *FLT3* indels, which appeared in diverse forms including seven cases of DUPs (32-73bp) and one case of small indel (30bp) (Supplementary Fig. 13). Taken together, these results indicate that ETCHING shows high performance for detecting SVs and FGs in both WGS and targeted sequencing data, indicating its general usability.

### Benchmarking computational efficiency

ETCHING significantly reduced the running time through implementation of the Filter module, resulting in computational speeds that were at least 15 times faster than those of the other tools (Fig. 1f). Such fast predictions result from significantly reduced genome mapping of reads. Although novoBreak also takes advantage of the *k*-mer approach to assembly TS contigs with BPs, it requires prior genome mapping of all reads to find read clusters, which is a time-consuming step. To confirm this conclusion, we determined the running times of ETCHING and novoBreak for each step (read filtration, mapping, and SV calling) on the HCC1395 dataset (Fig. 5a). As shown in Fig. 1f, based on its CPU time, ETCHING was approximately 15 times faster than novoBreak, mostly due to a reduction in the mapping time. In fact, most of novoBreak’s running time was spent in the mapping step (87%, 283.5 CPU-hours), whereas ETCHING used about 13% of its running time for this step (2.6 CPU-hours; Fig. 5a). Unlike other tools, ETCHING significantly reduces computational costs through its filtration-and-mapping strategy (Fig. 5b). Using multiple processes (30 threads) for parallel computing, ETCHING completed the entire procedure for nine hold-out datasets in 2.2h on average and for HCC1395 in 1.5h (Supplementary Fig. 5 and 14). Application of different numbers of threads showed that the efficiency of ETCHING approached saturation (1.5h) over 25 threads (Supplementary Fig. 15).

**Fig. 5.**
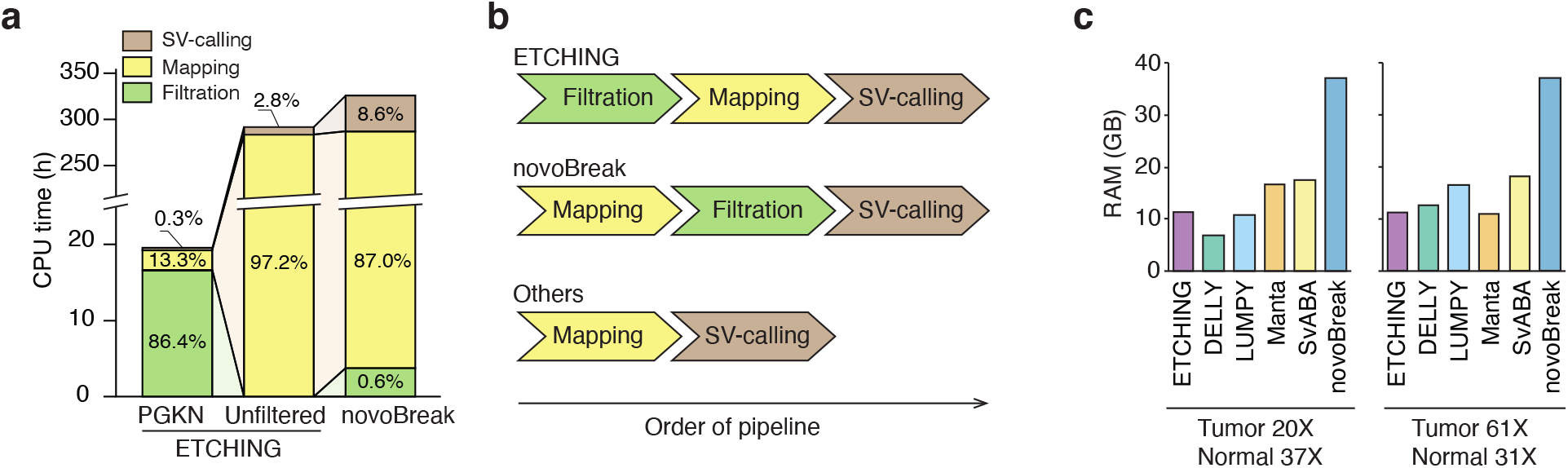
Computational costs of ETCHING and the benchmarking tools. **a.** Stepwise comparison of the CPU times for SV prediction using ETCHING, with reads filtered by PGKN or with unfiltered reads, and using novoBreak. **b.** Algorithmic differences between ETCHING, novoBreak, and others (DELLY, LUMPY, Manta, and SvABA). **c.** RAM usage by the SV callers on TCGA-A2-A04P (20X tumor, 37X normal) and TCGA-A1-A0SM (61X tumor, 31X normal) datasets with 60 threads.

Computational efficiency reflects both speed and memory usage, which have a trade-off relationship. However, benchmarking the memory usages of SV callers on 20X and 61X tumor samples showed no such relationship (Fig. 5c), which is probably because the memory usage is more dependent on the number of *k*-mers than the sequencing depth. In fact, ETCHING consistently used ~12G RAM, regardless of the size of the input dataset, which is comparable or more efficient than other tools in terms of memory usage. This fixed memory usage is mostly attributable to the size of the PGKN set, which is the least variable. Taken together, these results show that ETCHING is computationally very efficient, yet does not exhibit compromised performance.

## Discussion

Here, we introduced a high performing and very efficient SV caller, ETCHING, which takes advantage of a scalable PGK set (>3.9 × 10^9^ 31-mers). Matched normal samples can extend ETCHING to the PGKN *k*-mer set to enrich reads with somatic variations. *k*-mer counting, and searching for an exact *k*-mer in the large *k*-mer set, impose critical challenges on *k*-mer-based SV callers. ETCHING utilized K-mer Counter (KMC)^34^ for efficient *k*-mer counting and employed a parallel roll-encoding method for searching for TS *k*-mers, allowing a highly efficient *k*-mer processing method.

ETCHING has excellent potential for the prediction of somatic SVs, even without matched normal data. The PGK filter module can remove reads present in pan-genome or containing common variations from tumor sequencing data (Fig. 1b). Although ETCHING may produce FPs, it is still useful in the absence of matched normal data (Fig. 1g; Supplementary Fig. 6). This flexibility will be quite helpful, particularly for clinical sequencing, which often lacks such matched normal data (Fig. 4).

ETCHING found five additional druggable SV targets (in *ALK*, *NTRK1*, *BRCA2*, *PIK3CA*, and *AKT1*), three of which (*ALK*, *NTRK1*, and *BRCA2*) were validated by PCR analysis, in MM patients who did not carry SV biomarkers. *ALK* amplification is a potential molecular target in several cancers and *ALK* inhibitors could be beneficial to patients carrying such an *ALK* amplification ^35^. Because multiple SV events of DELs, DUPs, and INVs were detected around the *NTRK1* gene in SNUH19_MM01, the two most likely paths for their creation were confirmed by PCR (Figure 3e). Although the *NTRK1-LMNA* fusion is known to be a druggable target, the amplification of 1q23.1, where the *NTRK1* locus resides, has also been proposed as a candidate hotspot in the progression of MM ^36^. Because *BRCA2* loss of function is a known cancer driver, we examined biallelic inactivation of the *BRCA2* gene by searching for somatic or germline SNVs or indels at that locus but confirmed no clinically relevant variations in the other allele.

ETCHING can also predict other types of variations, such as germline and *de novo* mutations. With a *k*-mer set from a reference genome (such as hg19), it can predict germline SVs. If we use *k*-mers of parental genome sequences, ETCHING can find *de novo* mutations in offspring genomes. The current version of ETCHING predicts FG candidates from DNA sequencing data, but the detection of high-confidence FGs requires transcriptome data, such as RNA-seq. Such detection will be possible, without a need for other FG callers, by using a *k*-mer set of reference transcriptomes or RNA-seq data from normal samples. Hence, by the selection of an appropriate *k*-mer set, ETCHING can be a multi-purpose predictor for diverse types of genomic variations and FGs.

Although both ETCHING and novoBreak take advantage of TS reads to predict somatic SVs, the main strategy of ETCHING is distinct from that of novoBreak, which collects TS reads by comparing tumor and normal reads after mapping (the mapping-and-filtration approach). Instead, ETCHING uses a filtration-and-mapping approach, which makes ETCHING much faster than novoBreak, by as much as an order of magnitude (Fig. 5; Supplementary Fig. 15). In addition, novoBreak performs a local *de novo* assembly using the resulting TS reads to assemble TS contigs, which is another source of the heavy computational burden. The resulting contigs are aligned to a reference genome to predict SVs and BPs based on the mapping patterns of the contigs. Thus, the risk of misassembly also cannot be neglected. In contrast, ETCHING predicts all possible SVs using split-reads of TS reads and filters FPs by a RF module, achieving a low FP rate.

In summary, ETCHING is the fastest method for SV and FG prediction, and this speed has been achieved without compromising its performance or memory usages. We believe that our new approach will not only provide an efficient strategy for predicting various variations in mega-genome projects but will also contribute to real-time clinical applications.

## Data availability

WGS data from 26 MM samples can be downloaded from the Clinical & Omics Data Archive (CODA; registration number: R002594) of the Korean National Institute of Health. Targeted gene panel sequencing data from reference materials are available at our website (http://big.hanyang.ac.kr/ETCHING).

## Code availability

ETCHING was designed for 64-bit Linux systems. At least 16 GB of RAM is required. We recommend at least 64 GB. All source and binary codes used in the study are available at http://big.hanyang.ac.kr/ETCHING and GitHub (https://github.com/ETCHING-team/ETCHING).

## Acknowledgments

We thank all of the BIGLab members, and Professor Sun Kim of Seoul National University for critical reading and comments. This work was supported by the National Research Foundation (NRF) funded by the Ministry of Science & ICT (2014M3C9A3063541 to JWN) and by the Korean Health Technology R&D Project, Ministry of Health and Welfare, Republic of Korea (HI15C3224 to JWN).

## Ethics declarations

All MM samples used in this study were prepared under the Human Biospecimen Ethics Guidelines and were approved by the Internal Review Board (IRB) of SNUH.

## Competing interests

None declared

## METHODS

### *k*-mer counting

An efficient *k*-mer counting tool, KMC, was applied to count all possible *k*-mers (31-mers) from tumor and normal reads. *k*-mer counting can be done with multi-process (MP) computation. The results of *k*-mer counting are summarized in a histogram (Supplementary Fig. 1b) showing the *k*-mer depth (count) on the x-axis and the number of *k*-mers on the y-axis; the *k*-mer frequency shows a bimodal distribution for WGS data. A histogram of error-free *k*-mers is known to be close to a normal (Poisson) distribution, whereas rare *k*-mers, considered to be those with sequencing errors, show an exponentially decreasing curve over low depths. Hence, the local minimum was generally determined to be between *k*-mer depth 3–10, varying with the sequencing depth, quality, and tumor heterogeneity. Therefore, tumor *k*-mers with depth below the local minimum (the cutoff for erroneous *k*-mers) were removed, and the remaining error-free *k*-mers were subjected to the following steps (Supplementary Fig. 1a). For normal *k*-mers, those below *k*-mer depth 2 were removed and the remainder were added to the *k*-mer set (PGKN).

For targeted gene panel sequencing data, the local minimum is usually not presented as in WGS data. As the local minimum *k*-mer depth in WGS data is generally observed at a point about 10% of the distribution value at *k*-mer depth 2, we used the point as the local minimum *k*-mer depth in panel sequencing data.

### Roll-encoding

To efficiently process *k*-mers, we introduced a roll-encoding strategy, which encodes a *k*-mer to a series of 2-bit numbers by our encoding rules: A to 00, C to 01, G to 11, and T to 10. Because the *k*-1 nucleotides of the *i*-th and (*i+*1)th *k*-mers overlap, we can obtain the (*i+*1)th encoded *k*-mer simply by sliding a 2-bit number. This approach means that a new 2-bit number is added to the last nucleotide of the (*i+*1)th *k*-mer while the first 2-bit number is removed from the *i*-th encoded *k*-mer (Supplementary Fig. 16a). This procedure is repeated until the end of a read. Our roll-encoding also simultaneously encodes *k*-mer reverse complements. The smaller of the forward- and reverse-encoded values was stored as a canonical encoded *k*-mer. This roll-encoding method appeared to be faster than methods with conventionally encoded and ordinary (not encoded) *k*-mers (Supplementary Fig. 16b).

### The reference and normal *k*-mer sets

The reference *k*-mer set, PGK, is a unique set of *k*-mers from references (10 human genome assemblies; Supplementary Table 1) and those embedding common non-medical (nonpathogenic) SNPs in hg19 (GRCh37.p13) from dbSNP (release number 150). The normal *k*-mer set is from matched normal input reads. PGKN is a unique set of the PGK and the normal *k*-mers. For the YH_1.0 genome assembly, which includes uncertain bases, all possible nucleotides were assigned to generate the *k*-mer set. The reference *k*-mer set (PGK) is stored as a binary database file for reuse. The PGK binary file can be downloaded from our website (http://big.hanyang.ac.kr/ETCHING).

### Filter module

The saved reference *k*-mer set (PGK) is loaded to a hash table in the Filter module. If there is a matched normal sample as input, then normal *k*-mers are added to the *k*-mer set (PGK + Normal). When tumor sequencing data are used as the input, they are decomposed into tumor *k*-mers. The tumor *k*-mers are then searched in the reference *k*-mer sets (PGK or PGKN). The tumor *k*-mers present in the reference *k*-mer set are regarded as reference *k*-mers; otherwise, they are regarded as TS *k*-mers and subjected to the following read-collection step. The read-collection step collects TS reads embedding a TS *k*-mer. To speed up the read-collection step, a multi-processing procedure for simultaneously treating reading, collecting, and writing substeps was implemented (Supplementary Fig. 16c, d).

### Reduced read mapping

From the total input tumor reads, only TS reads collected through the Filter were mapped to the reference genome (hg19) using BWA-MEM with default parameters. We also used default parameters in read mapping for benchmarking tools.

### Caller module

After the TS reads are mapped, the Caller module finds BND candidates (BP pairs) by analyzing split reads with supplementary alignment (SA) tags, as follows (Supplementary Fig. 3). We focused on simply clipped pairs only, not on complex or double clipped reads, to reduce FP calls (Supplementary Fig. 3a). First, we defined a BP by its vector or chromosome (or contig/scaffold) name, its clipped position on the chromosome, and its clipped direction (Supplementary Fig. 3b). If a read was clipped in a region that is downstream of the BP, its clipped direction *s* is denoted as “+”. If a read was clipped in a region that is upstream of the BP, its clipped direction *s* is denoted as “-”. Thus, reads clipped at a locus can define a BND with a BP pair. A lack of SA tags in a clipped read indicates that there is a single BP that we called as a single-breakend (SND). Once all of the BNDs and SNDs are defined, BNDs are then classified by SV type (such as DEL, DUP, INV, or TRA), with their chromosome, BP position, and clipped direction information (Supplementary Fig. 3c).

### Sorter module

The Sorter module is a machine learning classifier that removes FP SVs from the Caller module outputs. Because ensemble machines usually show optimal performance in diverse problems, we applied RF (https://github.com/crflynn/skranger), and extreme gradient boosting (XGB, https://github.com/dmlc/xgboost) models to this study. To train the models, we randomly selected 31 training and 9 hold-out test samples from 55 BRCA samples (Supplementary Table 2; Supplementary Fig. 4a) as follows. We first predicted all possible SVs using five benchmarking SV callers and summarized tumor purities and sequencing depths for all 55 samples. Based on this information, we excluded (1) nine samples that had a low number of predicted SVs (<100) for at least one caller, (2) four samples with too many predicted SVs (>50,000 on average), and (3) two additional samples, one with the highest tumor read depth (93X) and one with the lowest tumor purity (0.474), to avoid extreme cases. From the remaining 40 samples, we randomly selected 31 and 9 samples so that there would be about a 3:1 ratio of SV candidates in the training and hold-out test datasets, respectively (Supplementary Fig. 4a). There were 894,333 and 278,627 SV candidates in 31 training and 9 hold-out samples. For training data, we selected 315,949 SV candidates detected by the Caller module, which were subjected to the training step of the Sorter module.

There is no ground truth exhaustively validated by experiments for the TCGA dataset. Thus, we used silver standard SVs detected by multiple SV callers. Of 315,949 SV candidates predicted by the Caller module in 31 training samples, 10,736 SVs were simultaneously predicted by at least three SV callers (Supplementary Fig. 4b). We regarded them as silver standard SVs and the remainder (314,507 SVs) as false (Supplementary Fig. 4b, c). With the true and false SVs, we trained the models with six different features – clipped-read count (*CR*), split-reads count (*SR*), supporting paired-end read count (*PE*), average mapping quality (*MQ*), depth difference (*DD*), and total length of clipped bases (*TC*) (see Supplementary Note for more details). Our training procedure consists of an outer 10-fold cross-validation (CV) loop for training and an inner 10-fold CV loop for model selection (Supplementary Fig. 4d). The SVs in the training samples were split evenly into eleven sets, including ten outer-training sets (TR_out) and one validation set (VA). During the outer 10-fold CV, a test set is selected (TE) from TR_out, and the remaining nine sets were subjected to inner-training (TR_in). The model selection process was done by inner 10-fold CV using TR_in, which was evaluated on TE. The procedure was iteratively performed through an outer 10-fold CV loop. A final model was obtained by averaging ten trained models. We validated the final model on the VA.

We then searched the optimal classification cutoffs of RF and XGB scores using the VA set (Supplementary Fig. 4f). F1-scores of RF (or XGB) showed robust performances in the range from 0.2 to 0.8 (from 0.05 to 0.95 for XGB). We used RF as default ML module in this study.

### Parameter optimization for benchmarking SV callers

ETCHING was benchmarked to the popular, high performing SV callers DELLY, LUMPY, Manta, SvABA, and novoBreak over WGS data, cfDNA reference materials, and targeted gene panel sequencing data from tumor samples. We also benchmarked BreaKmer for cfDNA reference materials and targeted gene panel sequencing data.

For a fair comparison on WGS datasets, we searched optimal parameters of benchmarking tools corresponding to the nearest points to the perfect performance (where precision and recall rates are 100%) over PR-curves on HCC1395 data (Fig. 2a). The point minimizes the distance, 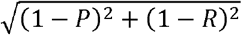, to (1,1) on given PR-curve, where *P* and *R* refer to precision and recall, respectively. DELLY’s optimal parameter was near its default parameter (−a 0.2), LUMPY was -m 12 option, and Manta was *minEdgeObservations* = 12 and *minCandidateSpanningCount* = 12. For SvABA, log-odd ratios of real and artifact variants ≥ 32 was the optimal one. novoBreak’s PR curve was closest to the corner for its statistical quality score ≥ 40. The statistical quality score is defined as 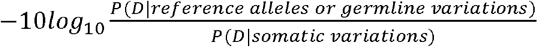, where *D* is the number of read counts supporting each variation or reference allele.

For cfDNA reference materials and targeted gene panel sequencing data, all tools were applied with default parameter sets. Manta was run with --tumorBam --exome options.

### Evaluation metrics

⍰ True positive (TP): Predicting true SVs (or biomarkers) as positive.
⍰ False negative (FN): Predicting true SVs (or biomarkers) as negative.
⍰ False positive (FP): Predicting false SVs (or biomarkers) as positive.
⍰ True negative (TN): Predicting false SVs (or biomarkers) as negative.

Given the TP, FN, FP, and TN metrics, the recall, sensitivity, precision, specificity, F1-score, and accuracy are estimated as follows:

- 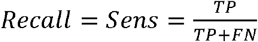
- 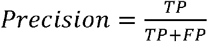
- 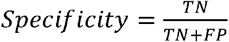
- 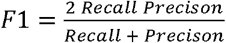
- 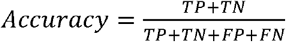

### Public WGS datasets

55 BRCA WGS datasets and 4 THCA WGS datasets were downloaded from TCGA (https://cancergenome.nih.gov).

### MM WGS data

Tumor cells were collected from bone marrow using CD138+ MACS sorting (Miltenyi Biotec, Auburn, CA) and DNA was extracted from the tumor cells for WGS library preparation. For matched normal samples, DNA was extracted from patients’ saliva with RNase treatment. Sequencing libraries were generated using a TruSeq nano DNA library prep kit (Illumina, San Diego, CA) following the manufacturer’s recommendations and sheared DNA fragments were end-repaired and size-selected to obtain DNA fragments around 350bp. Following PCR amplification, the DNA libraries were sequenced using the HiSeq™ X platform (Illumina). The 26 MM WGS datasets were produced and deposited in the CODA (registration number: R002594) of the Korean National Institute of Health. The study was approved by the Internal Review Board of Seoul National University Hospital (H-1103-004-353).

### FISH and karyotyping

Cytogenetic studies were performed at SNUH. Unstimulated bone marrow cells obtained at MM diagnosis were cultured for 24h; then, karyotypes were analyzed using the standard G-banding technique. The karyotypes were constructed and chromosomal abnormalities were reported according to the International System for Human Cytogenetic Nomenclature^37^. Interphase FISH was performed on myeloma cells from the bone marrow samples obtained at diagnosis according to the probe manufacturer’s instructions. Seven commercially available FISH probe sets were used. These included *IGH* dual-color, break-apart rearrangement probe; *TP53* SpectrumOrange probe; *RB1* D13S25 (13q14.3) SpectrumOrange probe; *IGH-FGFR3* dual-color, dual-fusion translocation probe; 1q21 SpectrumGreen probe; and *p16* (9p21, *CDKN2A*), SpectrumOrange/CEP9 SpectrumGreen probe (Abbott Diagnostics, Abbott Park, IL). The FISH experiments were performed on 26 MM specimens. The FISH probe sequences are summarized in Supplementary Table 6.

### PCR validation of actionable targets

PCR amplification was performed using the primer sets listed in Supplementary Table 9. Targets were amplified using primers designed in the flanking region of the junction. GAPDH was used as a control for assessing the PCR efficiency and for subsequent analysis by agarose gel electrophoresis.

### Manual curation of biomarkers and actionable targets in MM samples

The SV biomarkers and actionable targets were manually curated with all mapped reads. The candidate DELs and DUPs were checked by considering minor allele frequencies and read depth changes across chromosomes (Supplementary Fig. 11a,b), remaining focal DELs and DUPs. For *IGH*-associated TRA, candidate TRAs with which >10 paired-reads (mapping quality ≥20) are connected between *IGH* locus (14q32) and other loci in tumor but not in normal were selected as true somatic TRAs (Supplementary Fig. 11c,e). The candidates with the connection both in tumor and normal were considered as germline TRAs (Supplementary Fig. 11d). The read depth, minor allele frequency, discordant paired-read data to inspect true SVs during manual curation were summarized in Supplementary Material.

### Cell-free DNA reference materials

Targeted sequencing data from cfDNA reference materials (SeraCare, Milford, MA) were generated. DNA libraries were prepared using a KAPA Hyper Prep kit (Kapa Biosystems, Woburn, MA) as described previously. Hybrid selection for target enrichment was performed using customized baits targeting 38 cancer-related genes. After hybrid selection, the libraries were pooled, amplified, purified, quantified, and then subjected to cluster amplification according to the manufacturer’s protocol (Illumina). Flow cells were sequenced in the 150bp paired-end mode using a NextSeq 500/550 High Output Kit v2.5 (Illumina). The mean target coverage was 2023X. Two kinds of DNA mixtures, with the frequency of variant alleles ranging from 0.5–5.0% (CMM and MMv2), and a plasma-like DNA mixture, with the frequency of variant alleles ranging from 0.5–2.5% (CR), were generated along with WT DNA (Supplementary Table 10). The WT material was used as the matched normal. Note that DELLY displayed a low recall in normal-matched case, since it excessively removed SV calls using matched-normal data in the filtration step (Supplementary Fig. 12). The BreaKmer tool was excluded from this analysis because it failed to call variants from any sample, presumably because its approach is not feasible for such low allele frequencies.

### Cancer panel datasets

For cancer panel data of BreaKmer, we downloaded hybrid capture targeted gene panel data (110 replicates from 38 cancer samples). Because the normal samples that were provided are not matched to the cancer samples, they were excluded from the analysis. One sample with three replicates was also excluded from this analysis, since it was marked as non-cancer sample rather than diagnosed cancer type (SRR1304190-2). Two datasets (SRR1304204, SRR1304210) failed to run in at least one benchmarking tool, so the remaining 105 replicates from 37 sample (216X mean coverage of the targets) were analyzed. Because the sample labels in SRA are inconsistent with those in BreaKmer paper, we used ones described in the paper.

To reproduce the results of previous BreaKmer study, we needed to install the same version of BreaKmer with detailed information of target bait. However, we failed to install the same version of BreaKmer in their publication, and the bait information was also unavailable. Hence, we tested two other releases, v0.0.4 and v0.0.6. The version v0.0.6 found 78 true SVs out of 105 and only 487 additional calls, while v0.0.4 found 70 true SVs with 17,738 additional calls. Thus, we selected v0.0.6 for comparison. To substitute the missing target bait information, we used the genomic coordinates of target gene regions.

In case of novoBreak, it requires normal sequencing data. However, there is no matched normal samples in the panel data. For the reasons, we simulated WGS reads (30X coverage) from hg19 using an in-house script for novoBreak.

*FLT3* indel (30–73bp) and *KIT* deletion (48bp) were included in the list of known target alterations. As SvABA separately reports indels as output, we used SvABA high-confidence indel report along with its SVs. However, although Manta also reports indels, we did not use them because they are unfiltered candidates.

## Supplementary information

### Supplementary Note

Graph theory presentation for SV analysis

Six features for machine learning

Commands for benchmarking tools

High quality SVs

**Supplementary Fig. 1. a.** Detailed workflow of the ETCHING pipeline. MP and SP indicate multiple and single processing, respectively. **b.** A representative *k*-mer distribution of WGS data.

**Supplementary Fig. 2. a.** The size of the unique set containing the hg19 and PGK *k*-mers. **b.** The size of the unique set containing the *k*-mers not present in hg19.

**Supplementary Fig. 3. a.** The Caller module uses simply clipped reads (left side) but excludes reads that make complex clipped pairs (right side). **b.** Each BND is a pair of BPs, *i.e.* (BP_*i*_, BP_*j*_). An SND is a single-BND consisting of one BP with a dangling point, *i.e.* (BP_*i*_, ϕ). If a BP displays reads clipped in a direction at position *x* on chromosome *c*, we define that BP as a node (*c, x, s*), where *s* indicates its clipped direction (+1 or −1). **c.** Classification of SV types. For a BND (BP_*i*_, BP_*j*_), *x_i_* and *x_j_* are the positions on chromosome *c*, and *s_i_* and *s_j_* are the clipped directions of each BP. The table on the right side shows the classification criteria for SVs.

**Supplementary Fig. 4. a.** A flowchart for selecting training data from BRCA samples. **b.** A Venn diagram of SVs predicted by ETCHING and other tools in the training set. **c.** The numbers of SVs predicted by multiple callers were tallied in a histogram. The number of SVs are indicated on the y-axis and the number of tools that predicted the corresponding SVs are indicated on the x-axis. The vertical line denotes the cutoff for selecting silver standard SVs. **d.** A schematic workflow for training machine learning modules. **e.** Training and validation results of machine learning. **f.** Optimized cutoffs of machine learning methods. We set the optimized cutoff to 0.4 for RF and XGB.

**Supplementary Fig. 5.** The wall-clock times used by ETCHING and other tools on nine hold-out BRCA samples, which were measured using 30 threads.

**Supplementary Fig. 6.** The effectiveness of the Filter module on HCC1395 data. **a.** The percentage of TS reads that passed the PGK, Normal, and PGKN filters. **b.** The precision, recall, and F1-scores of the ETCHING results from total reads (unfiltered) and TS reads collected by the PGK, Normal, and PGKN filters.

**Supplementary Fig. 7.** Benchmarking results on THCA samples by SV type.

**Supplementary Fig. 8.** Strategies for HQ SV detection using tumor (HCC1395) and normal sequencing data (HCC1395 BL). **a**. The landscape of the depth change within the HQ DELs and HQ DUPs. **b**. The landscape of the discordant read-pairs connecting BPs within the HQ INVs and HQ TRAs. (**c** and **d**). The ROC curve (left) and the density distribution (right) for setting the cutoff using the depth difference score (DS) of DELs (**c**) and DUPs (**d**). **e**. The ROC curve for setting the cutoff using the connected-pair score (CS) of the HQ INVs and TRAs. **f**. Bar plot showing the count of HQ SVs and all SVs. DS and CS are defined in Supplementary Note.

**Supplementary Fig. 9.** Benchmarking results on BRCA, MM, and THCA samples with HQ SVs.

**Supplementary Fig. 10.** Benchmarking results on MM samples by SV type.

**Supplementary Fig. 11.** Rational for SV manual curation **a.***RB1* biomarker shown with unbalanced minor allele frequency (MAF) and read depth change on Chr13 in MM20. **b.***AKT1* DUP locus shown with unbalanced MAF and read depth change on Chr14 in MM22. **c.** Discordant paired-reads connected between *IGH* locus and Chr11 of MM6. The blue dot near 69M indicates a TRA, t(11;14)(q13;q32), including *CCND1* gene in tumor. **d.** Germline TRA with discordant paired-reads both in tumor and normal. **e.** An instance of manually curated TRAs is shown in IGV.

**Supplementary Fig. 12.** SV and FG prediction on targeted gene panel sequencing data paired with sequencing data from WT alleles (regarded as matched normal). Otherwise, as in Fig. 4a–c.

**Supplementary Fig. 13.** Indels associated with *FLT3* in eight different samples. The index numbers are the same ones in Fig. 4d.

**Supplementary Fig. 14.** The wall-clock times used by ETCHING, ETCHING without filter (unfiltered), and novoBreak on HCC1395 data on 30 threads.

**Supplementary Fig. 15.** The wall-clock times used by ETCHING with different thread numbers ranging from 5 to 50.

**Supplementary Fig. 16. a.** Schematic of the roll-encoding algorithm for processing *k*-mers. As a *k*-mer window slides, it updates an encoded value using our encoding rule. **b.** The computing costs of the Read-collector using the conventional encoding method, ordinary *k*-mers, and roll-encoding methods on tumor (46X) and normal (31X) WGS data with 30 threads. **c.** A schematic workflow of parallel computing for read collection. **d.** Data from the read collection step are processed by parallel computing.

### Supplementary Tables

**Supplementary Table 1.** Reference genomes and dbSNP used in PGK

**Supplementary Table 2.** BRCA samples

**Supplementary Table 3.** THCA samples

**Supplementary Table 4.** MM samples

**Supplementary Table 5.** FISH and karyotype in MM samples

**Supplementary Table 6.** FISH probe sets

**Supplementary Table 7.** Manually curated partner BPs of *IGH* TRAs.

**Supplementary Table 8.** BreaKmer panel data

**Supplementary Table 9.** PCR primer sets

**Supplementary Table 10.** Reference material

## Notes

http://big.hanyang.ac.kr/ETCHING/

### Competing Interest Statement

The authors have declared no competing interest.

