## Supplementary figures and images for "Ultra-fast Prediction of Somatic Structural Variations by Reduced Read Mapping via Pan-Genome *k*-mer Sets"

### SuppFigure1.pdf

##### Supplementary Fig. 1

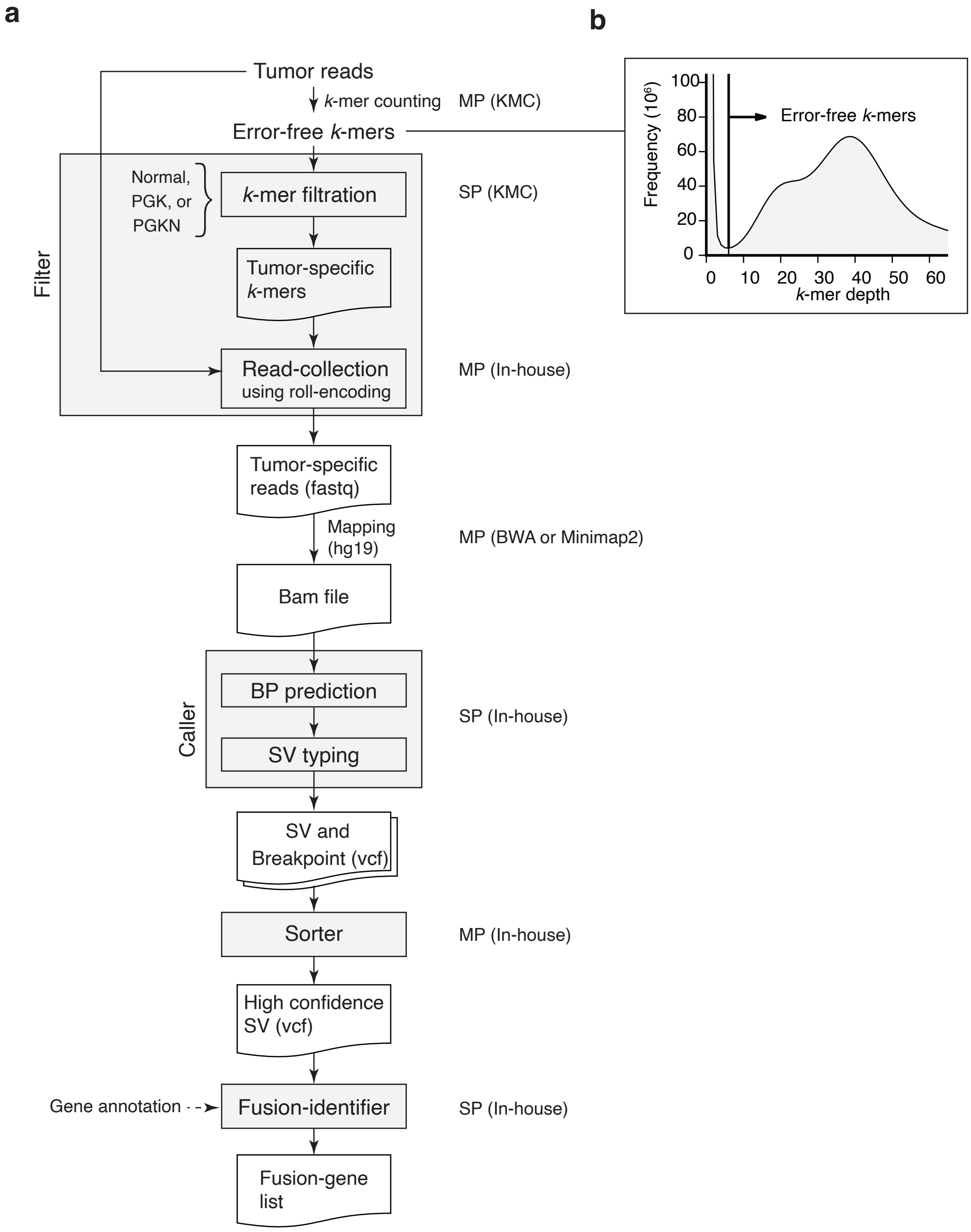

### SuppFigure2.pdf

##### Supplementary Fig. 2

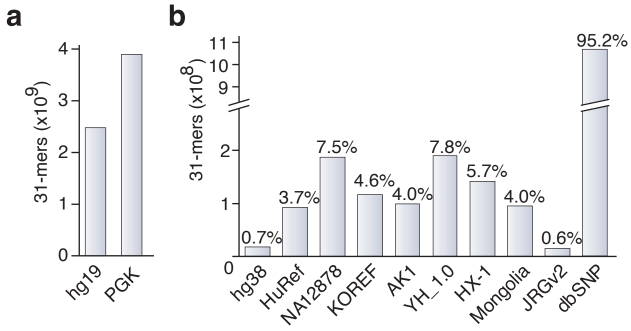

### SuppFigure3.pdf

#### Supplementary Fig. 3

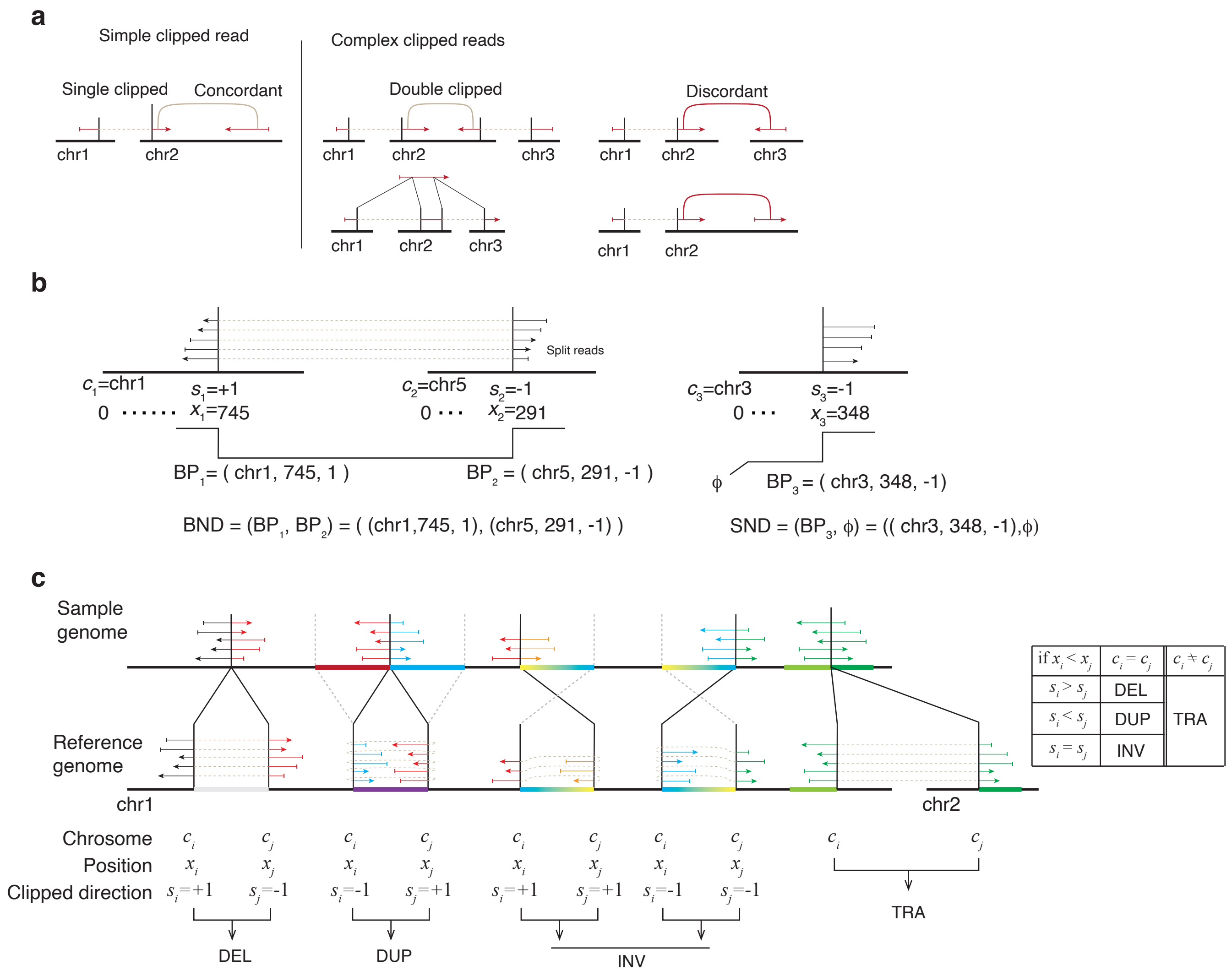

### SuppFigure4.pdf

#### Supplementary Fig. 4

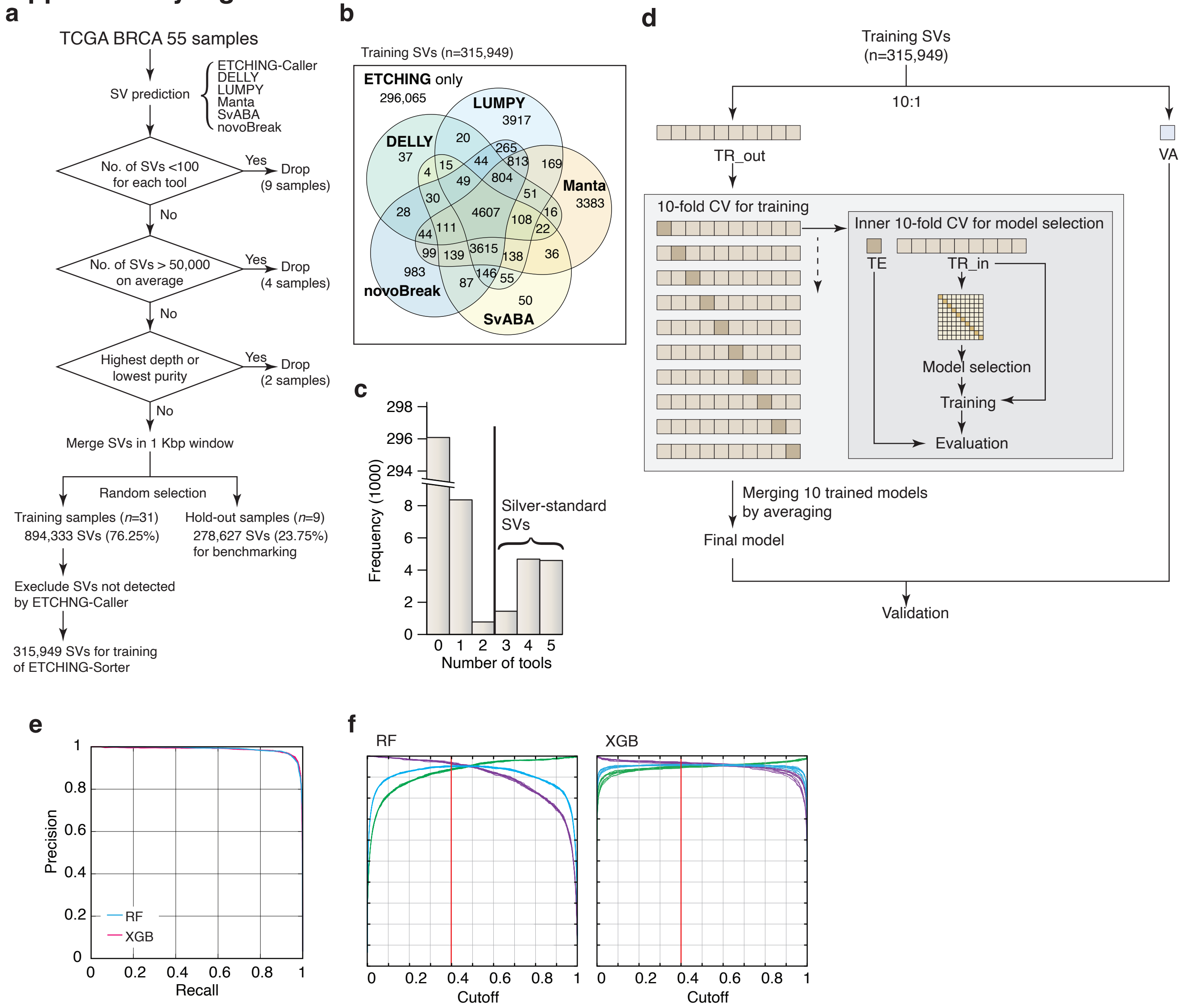

### SuppFigure5.pdf

**Supplementary Fig. 5**

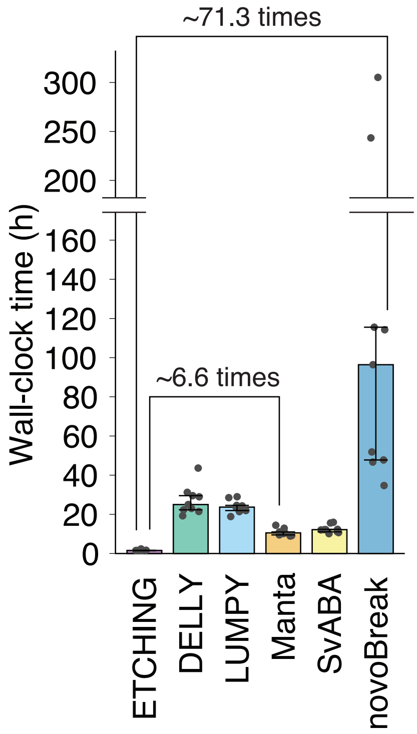

### SuppFigure6.pdf

##### Supplementary Fig. 6

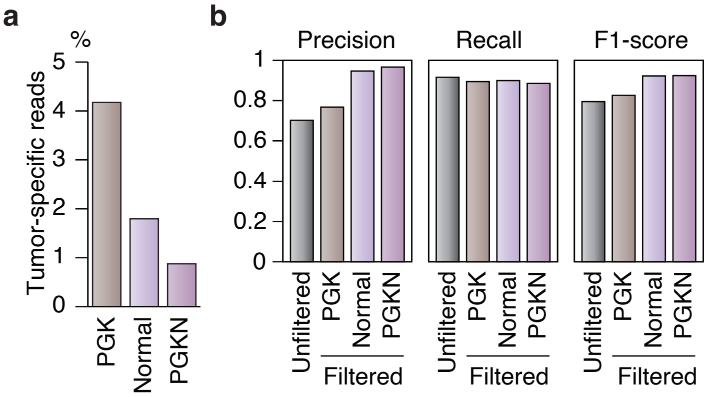

### SuppFigure7.pdf

#### Supplementary Fig. 7

##### Thyroid cancer

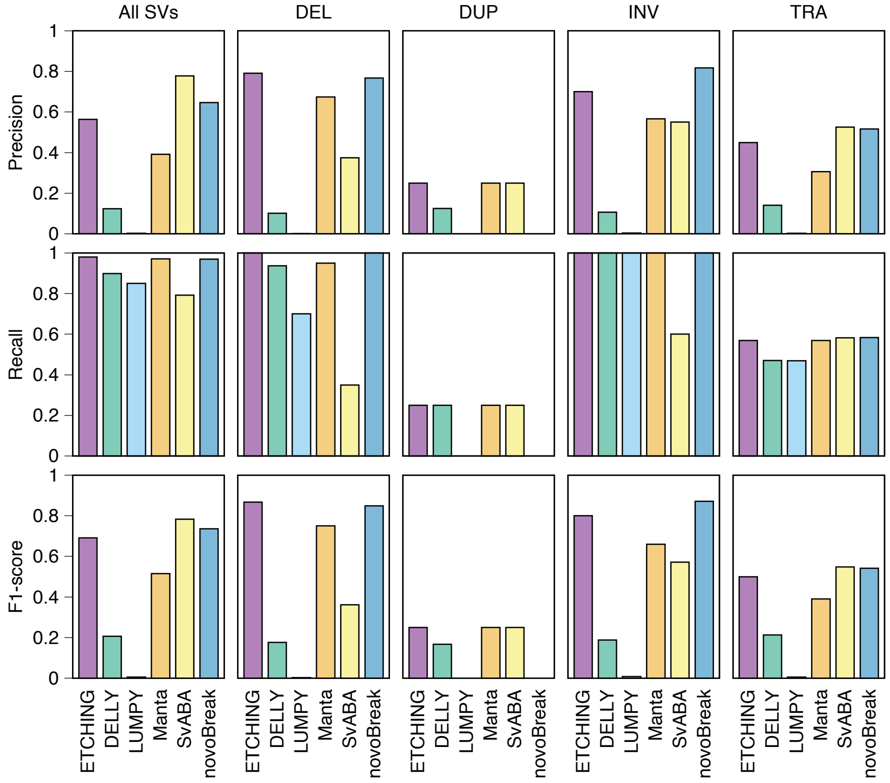

### SuppFigure8.pdf

#### Supplementary Fig. 8

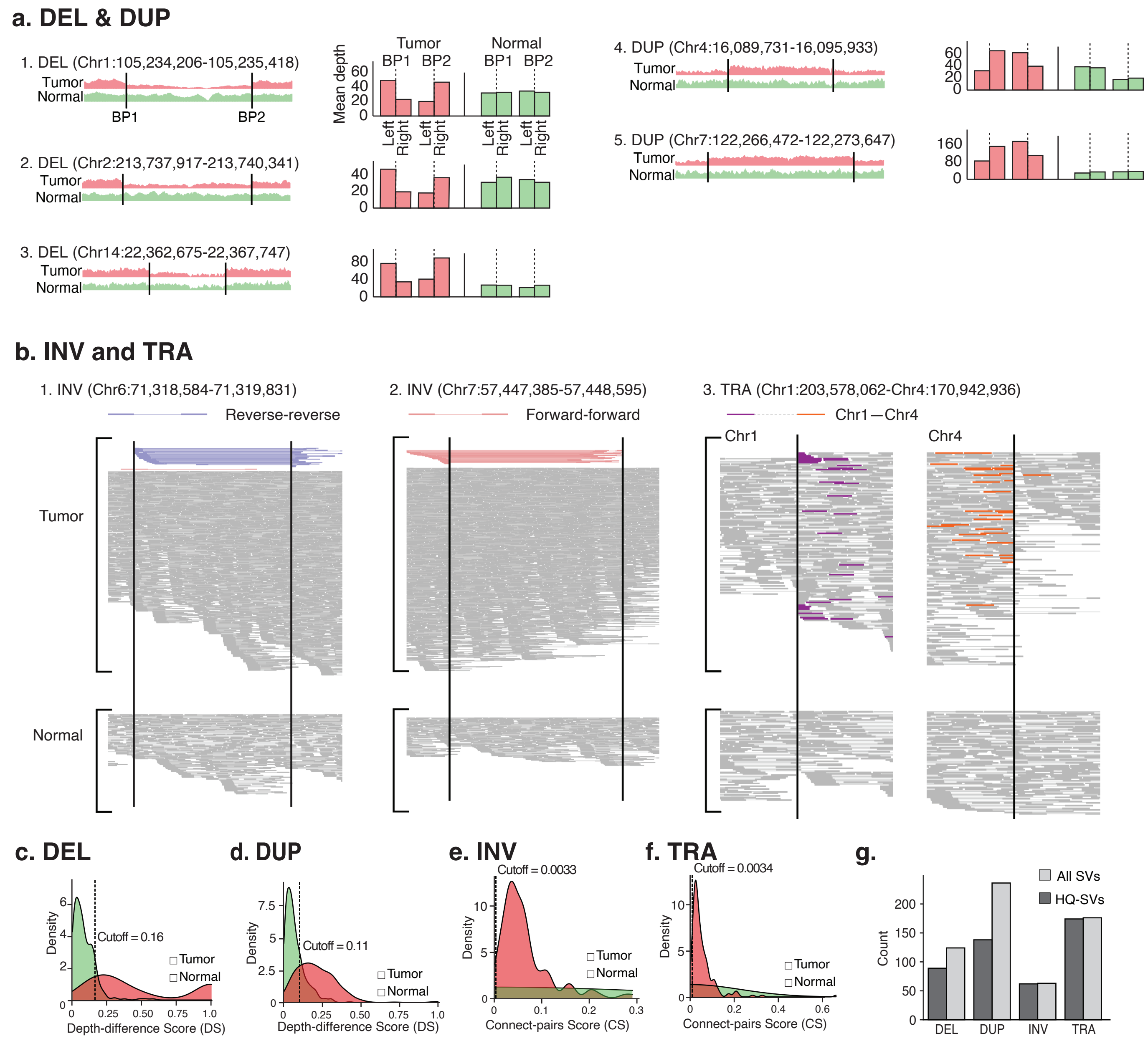

### SuppFigure9.pdf

#### Supplementary Fig. 9

##### HQ SVs

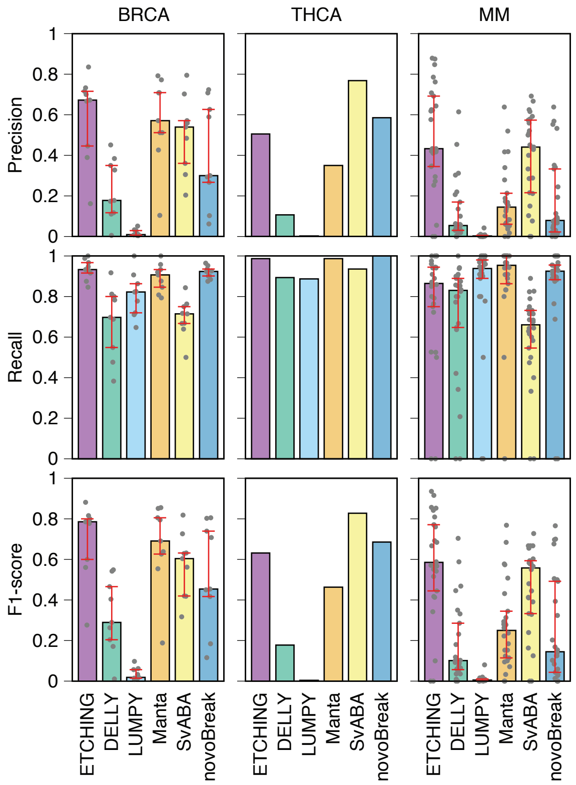

### SuppFigure10.pdf

##### Supplementary Fig. 10

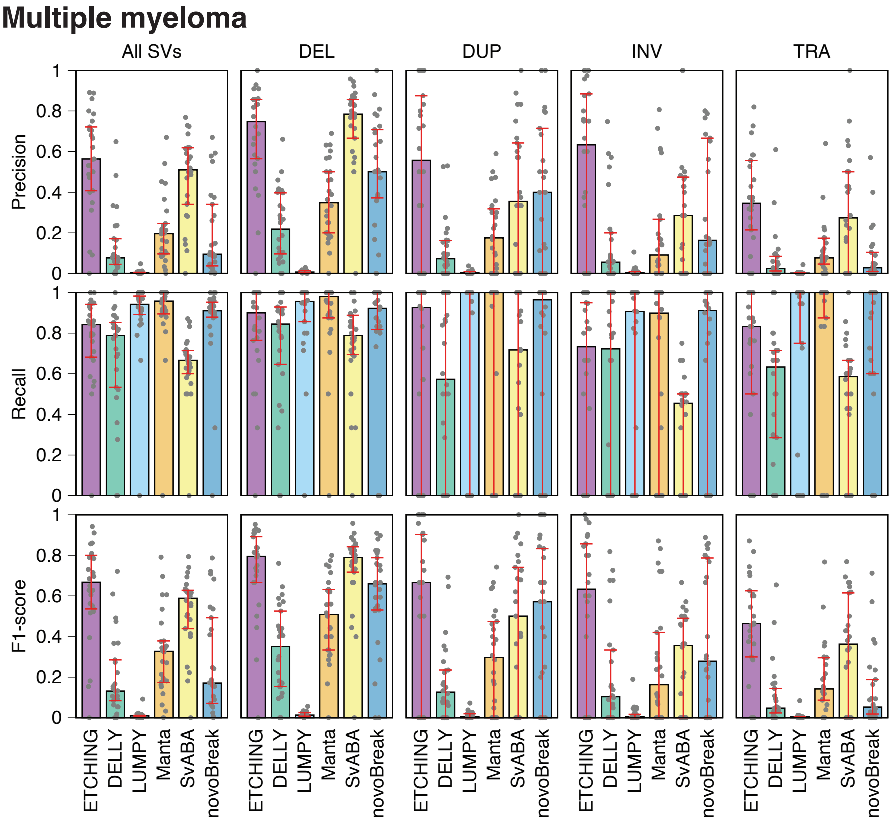

### SuppFigure11.pdf

##### Supplementary Fig. 11

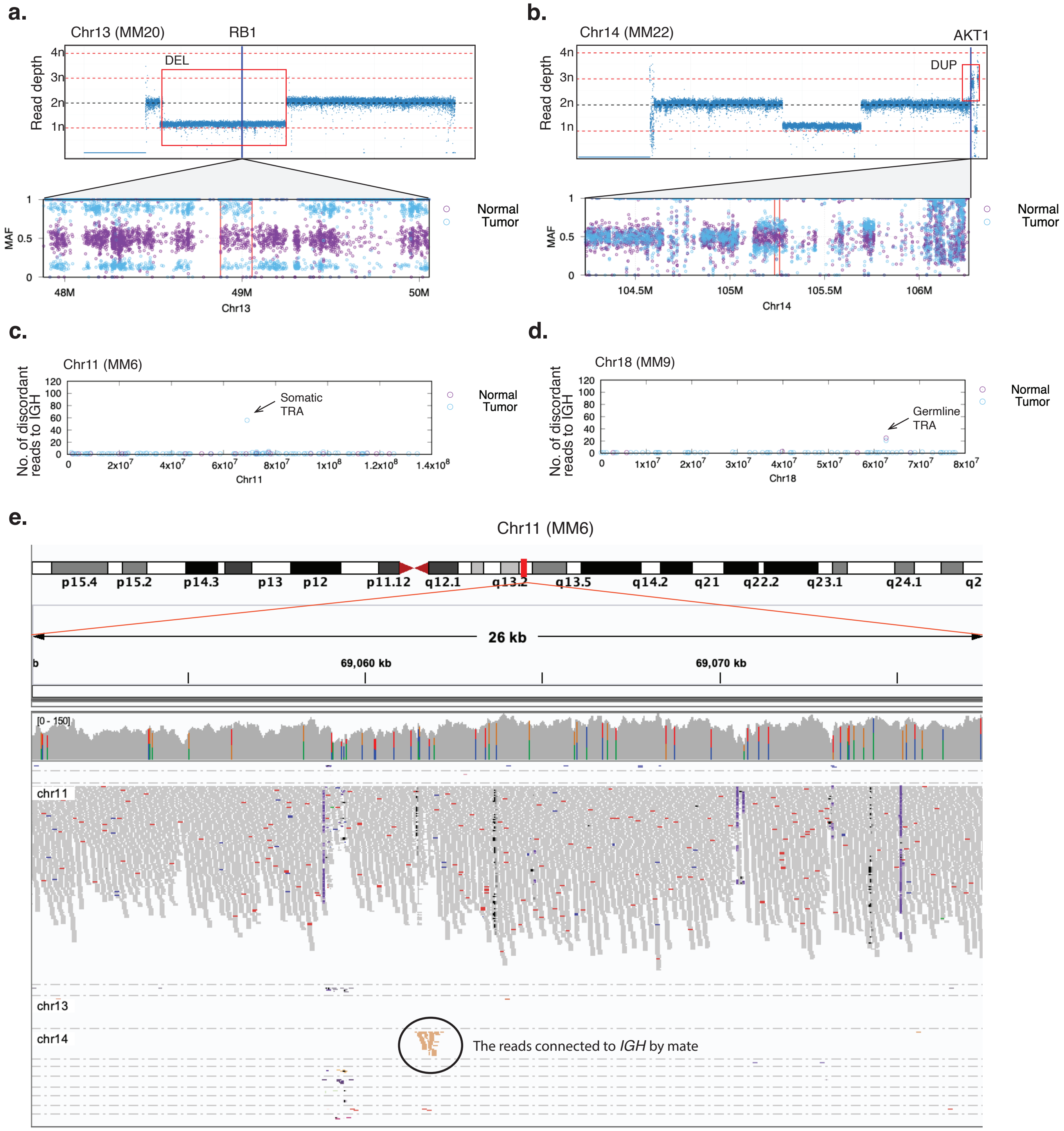

### SuppFigure12.pdf

##### Supplementary Fig. 12

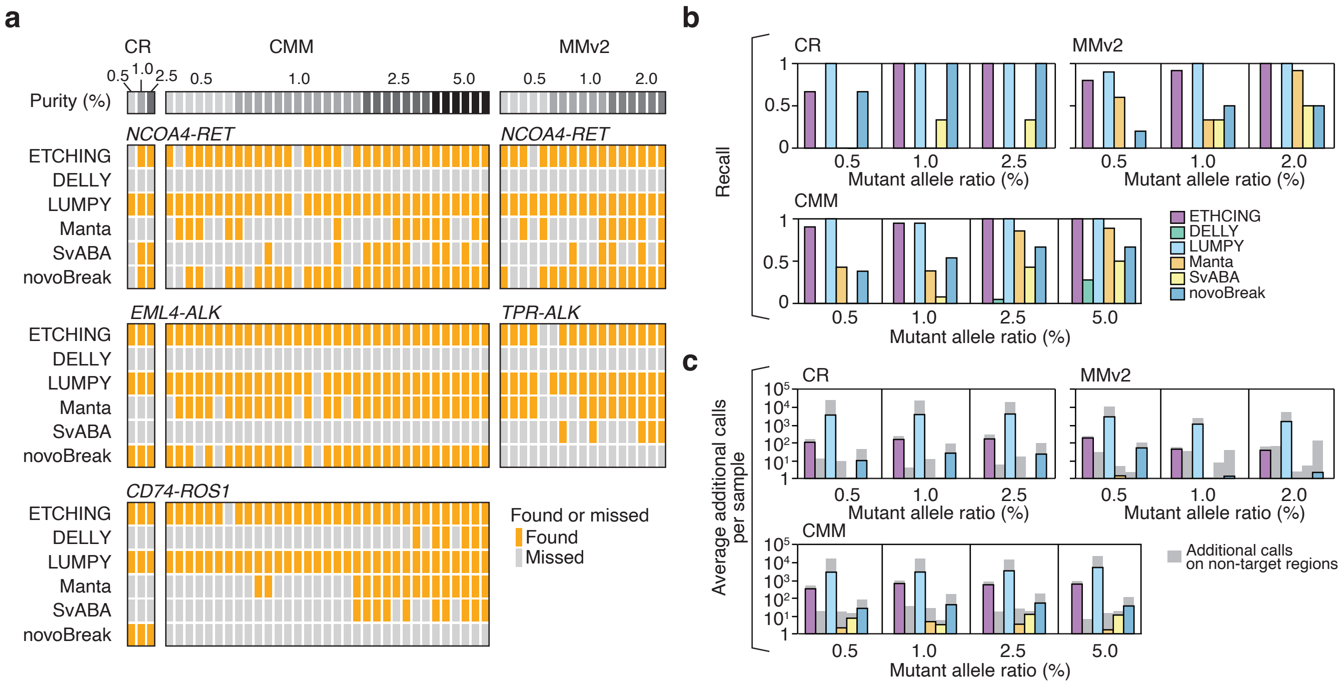

### SuppFigure13.pdf

##### Supplementary Fig. 13

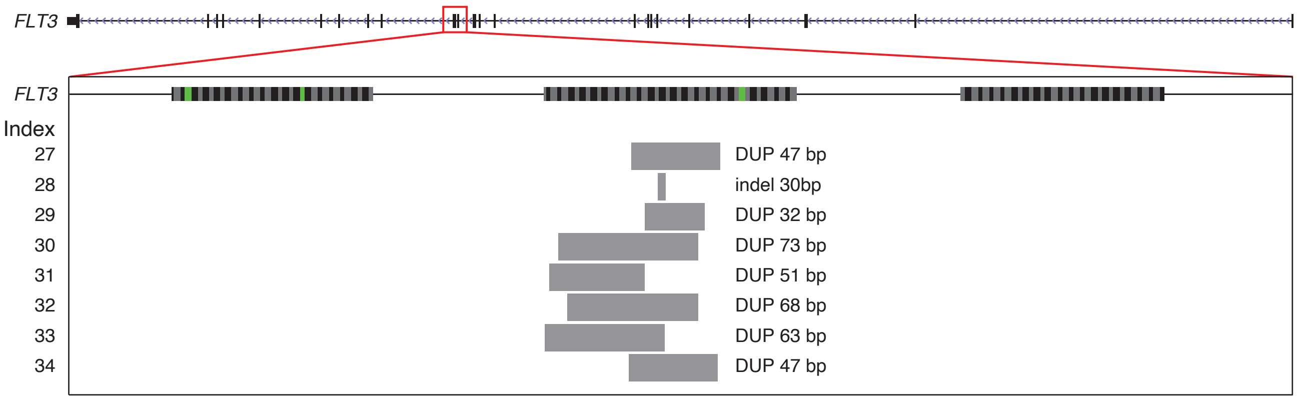

### SuppFigure14.pdf

##### Supplementary Fig. 14

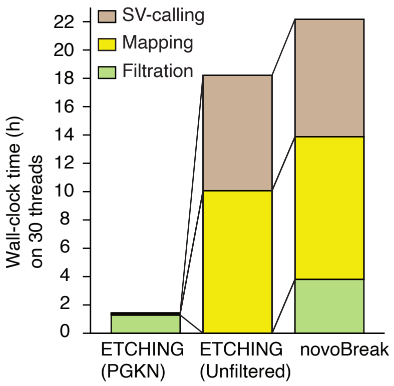

### SuppFigure15.pdf

##### Supplementary Fig. 15

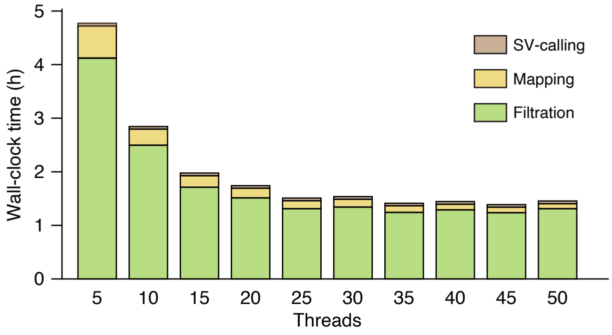

### SuppFigure16.pdf

##### Supplementary Fig. 16

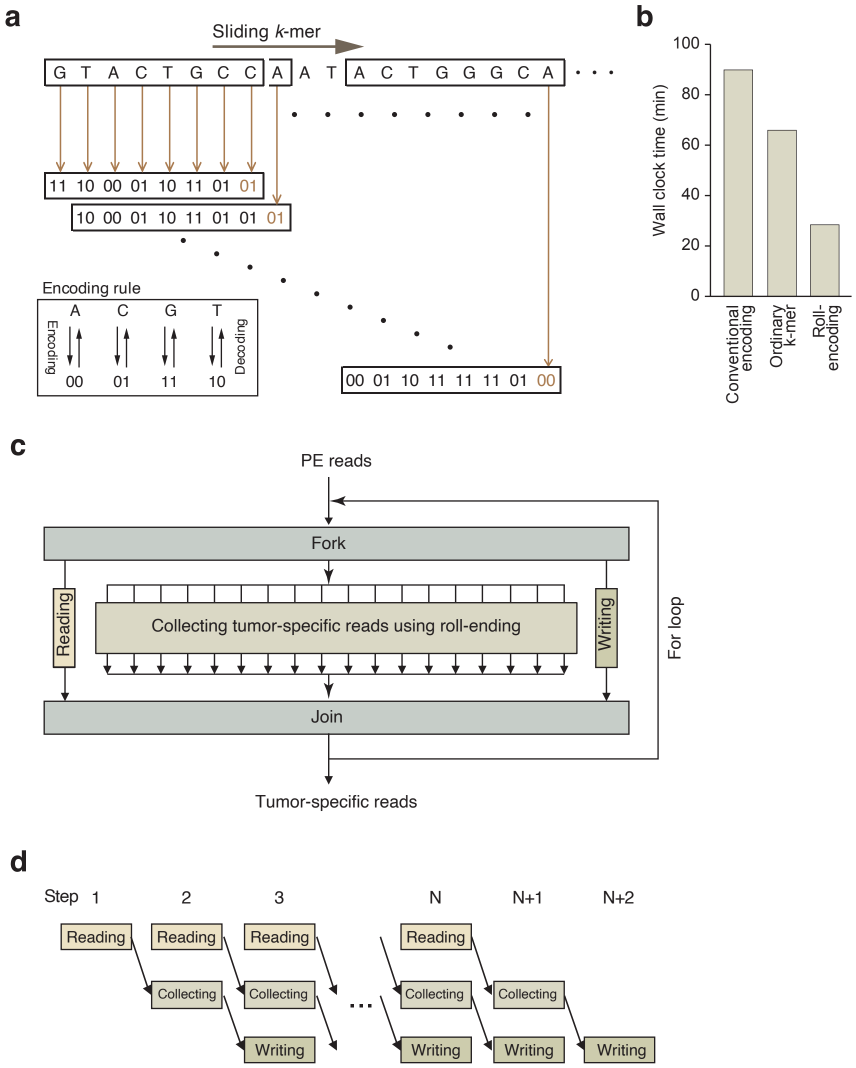
