## Supplementary Note for "Ultra-fast Prediction of Somatic Structural Variations by Reduced Read Mapping via Pan-Genome *k*-mer Sets"

**Graph theory presentation for SV analysis**

Let us suppose that there is a (reference) genome consisting of chromosomes, $C=\{c\}$, where c is a chromosome. In the case of the human genome,

$c\in\{chr1, chr2, \cdots,X,Y\}$.

Unplaced chromosomes can be included in the set. On a chromosome *c*, position and strand (or direction) can be written as

$x\in[1,L(c)]$, and

$s\in\{1,-1\}$,

respectively, where $L(c)$ is the size of chromosome *c*, and $\pm1$ indicates the forward/reverse (5’-to-3’/3’-to-5’) direction in *s*. Then, we can describe SVs using graph theory,

$\mathbf{G}_{SV}=(\mathbf{BP},\mathbf{BND})$,

where BP and BND refer to node and edge, respectively as follow.

$\mathbf{BP}=\left\{ BP | BP=(c,x,s) \right\}$, and

$\mathbf{BND}=\left\{ BND | \mathrm{BND}_{ij}=\left( \mathrm{BP}_{i},\mathrm{BP}_{j} \right) \mathrm{and} \mathrm{BP}_{i}≇\mathrm{BP}_{j} \right\}$.

$BP_{i}≇{BP}_{j}$ means that $c_{i}\neq c_{j}$, $\left| x_{i}-x_{j} \right|\geq l$, or $s_{i}\neq s_{j}$, where $l$ is the minimum size ($\geq1)$ of the SV. For a large SV, $l$ is generally 100 or 1000.

For instance, if a read(s) is clipped at genomic position 745 on chr1 in the (+) direction, the BP can be written as

$\mathrm{BP}_{1}=\left( chr1, 745,1 \right)$.

In the same way, if its mate BP is located at genomic position 271 on chr5 in the (-) direction, it can be denoted as

$\mathrm{BP}_{2}=\left( chr5, 271,-1 \right)$.

Then, $\mathrm{BND}_{12}=\left( \mathrm{BP}_{1},\mathrm{BP}_{2} \right)$ can be denoted as

$\mathrm{BND}_{12}=\left( \left( chr1, 745,1 \right), (chr5, 271,-1) \right)$.

Let us define the set of SVs comprising DEL, DUP, INV, and TRA, *i.e.*

$$\mathbf{SV}\boldsymbol{\equiv}\bigcup\mathbf{X}\boldsymbol{,}$$

where $\boldsymbol{X\in\{DEL,DUP,INV,TRA}\}$, and for $x_{i}<x_{j}$,

$\mathbf{DEL}=\left\{ \left( \mathrm{BP}_{i},\mathrm{BP}_{j} \right) | (s_{i}>s_{j})\wedge\left( c_{i}=c_{j} \right) \right\}$,

$\mathbf{DUP}=\left\{ \left( \mathrm{BP}_{i},\mathrm{BP}_{j} \right) | (s_{i}<s_{j})\wedge\left( c_{i}=c_{j} \right) \right\}$,

$\mathbf{INV}=\left\{ \left( \mathrm{BP}_{i},\mathrm{BP}_{j} \right) | (s_{i}=s_{j})\wedge\left( c_{i}=c_{j} \right) \right\}$, and

$$\mathbf{TRA}=\left\{ \left( \mathrm{BP}_{i},\mathrm{BP}_{j} \right) | c_{i}\neq c_{j} \right\}$$

(Supplementary Figure 3c).

As a particular case, single-breakend (SND) can be defined as,

$\mathbf{SND}=\left\{ \mathrm{SND}_{i}=\left( \mathrm{BP}_{i},\mathrm{BP}_{unknown} \right) \right\}⊈\mathbf{BND}$,

where $\mathrm{BP}_{unknown}\notin\mathbf{BP}$. Thus, an SND is a path to an unknown sequence, and it could be any type of SV including INS. It implies that an SND may be related to an INS event.

**Six features for machine learning**

The Sorter module uses six BND features, which are the clipped-read (*CR*) count, split-reads (*SR*) count, supporting paired-end (*PE*) read count, average mapping quality (*MQ*), depth difference (*DD*) in the upstream and downstream 100 bps of a BP, and total length of clipped bases (*TC*). *SR* and *PE* are BND features, and *CR*, *MQ*, *DD*, and *TC* are BP-derived BND features. These features were extracted from TS reads mapped to genomes.

The BND features, ${SR}_{ij}$ and ${PE}_{ij}$, are given by the number of split- and paired-end reads, respectively, supporting ${SV}_{ij}=({BP}_{i},{BP}_{j})$. BP-derived BND features, $F_{ij}$ (one of $CR$, $MQ$, $DD$, and $TC$), are calculated by harmonic averages ($\frac{2f_{i}f_{j}}{f_{i}+f_{j}}$) of BP features, $f_{i orj}$ (one of *cr*, *mq*, *dd*, and *tc*). Here, *cr* is the number of clipped-reads at a BP, *mq* is an averaged mapping quality of reads mapped in the range of read-lengths on both sides based on the BP, *dd* is the difference of averaged depths in the range of read-lengths of TS reads between both sides of the BP, and *tc* is the sum of clipped lengths at a BP. In the case of an SND, *SR* is given by the number of clipped reads among mate-unmapped reads, *PE* is given by the number of mate-unmapped reads, and all BP-derived SND features are given by themselves, *i.e.* $F=f$, because an SND consists of only one BP.

Because all of the features, with the exception of *MQ*, are proportional to the sequencing depth (*D*) and tumor purity (*P*), we normalized the BND features with *D* and *P*, i.e. $\frac{SR}{D\times P}\to SR$, and so on. *MQ* was used as-is because it is not related to *D* or *P*.

**Commands for benchmarking tools**

We ran benchmarking tools as follows.

***ETCHING-***To run ETCHING, a configuration file must be written as follows:

$ vi example.conf

sample	tumor_1.fastq

sample	tumor_2.fastq

filter_fq	normal_1.fastq

filter_fq	normal_2.fastq

filter_fa	assembled_genome.fasta

filter_db	PGK

genome	hg19.fa

annotation	hg19.gtf

Then, the ETCHING pipeline can be run by a simple command.

$ etching example.conf

In the pipeline, etching_filter, mapping, etching_caller, etching_sorter, and etching_fg_indendifier are performed sequentially. The pipeline can be run step-by-step for WGS data as follows.

$ etching_filter example.conf -p result

$ etching_caller -b result.sort.bam -g hg19.fa -o result

$ etching_sorter -i result.BND.vcf -o result.BND

or

$ etching_sorter -i result.SV.vcf -o result.SV

Optionally, FGs can be identified by a command,

$ etching_fg_identifier result.BND.etching_sorter.vcf annotation_gencode.gtf

*k*-mer filtration, read-collection, and mapping steps were included in etching_filter. Based on the mapping results of etching_filter, etching_caller finds BNDs or SVs. If total tumor and normal reads were used, somatic SVs can be selected by

$ somatic_filter tumor.vcf normal.vcf > somatic.vcf

For other benchmarking tools that do not determine the SV type, we prepared a simple SV-typer, which can be run by the command,

$ etching_typer input.vcf

***Alignment–***For other tools, we aligned normal and tumor sequencing data on the hg19 genome:

$ bwa mem -t 30 -R \"@RG\tID:tumor\tSM:tumor\tLB:WGS\" -V \

hg19.fa tumor_1.fq tumor_2.fq | samtools view -Sbh - > tumor.bam

$ bwa mem -t 30 -R \"@RG\tID:normal\tSM:normal\tLB:WGS\" -V \

hg19.fa normal_1.fq normal_2.fq | samtools view -Sbh - > normal.bam

Aligned sam files were compressed to bam files, which were sorted by coordination.

$ samtools sort tumor.bam -o tumor.sort.bam

$ samtools sort normal.bam -o normal.sort.bam.

***DELLY–***Using DELLY (version 0.7.9) ^1^, we made an initial vcf file, and filtered it:

$ delly call -x exclude.tsv -o Delly.bcf -g hg19.fa tumor.sort.bam normal.sort.bam

$ echo -e "tumor\ttumor\nnormal\tcontrol" > samples.tsv

$ delly filter -f somatic -o Delly.filter.bcf -s samples.tsv -a *value* Delly.bcf

***LUMPY–***LUMPY (version 0.3.0) ^2^ requires several preparation steps. For tumor sequencing data, we conducted preparation steps as follows:

$ samtools view -h tumor.bam \

| samblaster --excludeDups --addMateTags --maxSplitCount 2 --minNonOverlap 20 \

| samtools view -Sbh - > tumor.lumpy.bam

$ samtools view -b -F 1294 tumor.lumpy.bam > tumor.discordant.bam

$ samtools view -h tumor.lump| extractSplitReads_BwaMem -i stdin | samtools view -Sbh - \

> tumor.split.bam

$ samtools sort tumor.lumpy.bam -o tumor.lumpy.sort.bam

$ samtools sort tumor.discordant.bam -o tumor.discordant.sort.bam

$ samtools sort tumor.split.bam -o tumor.split.sort.bam

We repeated the same steps for normal sequencing data. Then, we ran LUMPY (lumpyexpress version 0.3.0) several times while changing the minimum sample weight for a call (option -m) as follows:

$ *weight*=12

$ lumpy/lumpyexpress \

-B tumor.lumpy.sort.bam,normal.lumpy.sort.bam \

-S tumor.split.sort.bam,normal.split.sort.bam \

-D tumor.discordants.sort.bam,normal.discordants.sort.bam \

-o lumpy_m${i}.vcf \

-m ${*weight*}

for different weights.

***Manta*–**To run Manta (version 1.6.0) ^3^, we modified a configuration file (manta.conf) as follows:

$ *manta_parameter*=12

$ echo "[manta]

*minEdgeObservations* = ${*manta_parameter*}

*minCandidateSpanningCount* = ${*manta_parameter*}

referenceFasta = ${genome}

minCandidateVariantSize = 8

rnaMinCandidateVariantSize = 1000

graphNodeMaxEdgeCount = 10

minScoredVariantSize = 50

minDiploidVariantScore = 10

minPassDiploidVariantScore = 20

minPassDiploidGTScore = 15

minSomaticScore = 10

minPassSomaticScore = 30

enableRemoteReadRetrievalForInsertionsInGermlineCallingModes = 1

enableRemoteReadRetrievalForInsertionsInCancerCallingModes = 0

useOverlapPairEvidence = 0" > manta.conf

Then, we ran the commands as follows:

$ python configManta.py \

--normalBam=normal.sort.bam --tumorBam=tumor.sort.bam \

--referenceFasta=hg19.fa --config=manta.conf --runDir=./

$ python Workflow.py -m local

***SvABA*–**Once we ran SvABA (version 1.1.3) ^3^ with default settings, we performed post-processing by removing the SVs with a high LOD value in the vcf file. The following command obtained the initial result:

$ svaba run -t tumor.sort.bam -n normal.sort.bam -D dbsnp.vcf -a initial -G hg19.fa

We filtered low-quality SVs not satisfying the condition *LO*>=*cut-off* from initial.svaba.somatic.vcf file. Here *LO* is *log-odds* value, which is recorded in the vcf file. SvABA’s F1-score maximized (0.901) when the *cut-off* was 32.

***novoBreak*–**Similarly, as we did for SvABA, we removed low-quality SVs from the initial result after running novoBreak (version 1.1) ^4^. The initial result was obtained by the following command:

$ run_novoBreak.sh novoBreak hg19.fa tumor.sort.bam normal.sort.bam \

30 working_directory

***Performance calculation*–**For all vcf files generated by the benchmarking tools, we applied our SV typing tool, etching_typer, to classify unclassified BNDs as DEL, DUP, INV, or TRA. In the case of DELLY and LUMPY, BNDs have no mate, whereas the others generate mates. Thus, we added their mates by parsing BND for DELLY and LUMPY. Then, we merged all vcf files from benchmarking tools (ETCHING, DELLY, LUMPY, Manta, SvABA, and novoBreak) using SURVIVOR ^5^:

$ echo ETCHING.vcf > benchmarking.list

$ echo DELLY.vcf >> benchmarking.list

$ echo LUMPY.vcf >> benchmarking.list

$ echo Manta.vcf >> benchmarking.list

$ echo SvABA.vcf >> benchmarking.list

$ echo novoBreak.vcf >> benchmarking.list

$ SURVIVOR merge benchmarking.list 10 1 1 1 0 1000 benchmarking.merge.vcf

Parsing the merged vcf file (benchmarking.merge.vcf), we measured precision, recall, and F1-score parsing format field of the merged vcf file. The format field is recorded in the order of benchmarking.list. If “./.:NaN:0:0,0:--:NaN:NaN:NaN:NAN:NAN:NAN” was recorded at *i*’th field in the merged vcf file, it means that the *i*’th tool did not find the SV; otherwise, the tool found the SV.

**High quality SVs**

Criteria for HQ SVs were based on analysis of HCC1395 data (Supplementary Fig. 8). Because DEL and DUP generally show significant depth differences, we introduced the depth difference score (DS) of a BP,

$DS=\frac{\left| D_{1}-D_{2} \right|}{D_{1}+D_{2}}$

for DEL and DUP, where $D_{1}$ and $D_{2}$ are average read depths in the range of 150 bp on the left and right sides of a BP, respectively. For an SV with BP_1_ and BP_2_, we calculated ${DS}_{1}^{t}$ and ${DS}_{2}^{t}$ for tumor data, and ${DS}_{1}^{n}$ and ${DS}_{2}^{n}$ for normal data. If

$\left( {DS}_{1}^{t} \mathrm{or} {DS}_{2}^{t} \right)\geq\mathrm{cutoff}$, and

$\left( {DS}_{1}^{n} \mathrm{and} {DS}_{2}^{n} \right)<\mathrm{cutoff}$,

we selected it as a HQ SV; otherwise, it was dropped. For INV and TRA, we used the number of discordant reads supporting an SV, because there is no depth difference in such SVs. Thus, for INV and TRA, we introduced a connection pair score (CS),

$CS=C/(P+C)$,

where *P* and *C* are the numbers of concordant and discordant pairs, respectively, within the range of $\pm$500 bp. In the same manner, as we did for DEL and DUP, we selected HQ INVs and TRAs, which satisfy

$\left( {CS}_{1}^{t} \mathrm{or} {CS}_{2}^{t} \right)\geq\mathrm{cutoff}$, and

$\left( {CS}_{1}^{n} \mathrm{and} {CS}_{2}^{n} \right)<\mathrm{cutoff}$.

Based on the ROC curves using the HQ SVs, we determined the cutoffs, which are 0.16 for DEL, 0.11 for DUP, 0.0033 for INV, and 0.0034 for TRA (Supplementary Fig. 8). Using HQ SVs, we calculated performances.
